# scAlign: a tool for alignment, integration and rare cell identification from scRNA-seq data

**DOI:** 10.1101/504944

**Authors:** Nelson Johansen, Gerald Quon

**Affiliations:** Graduate Group in Computer Science, University of California, Davis, Davis, CA; Genome Center, University of California, Davis, Davis, CA; Department of Molecular and Cellular Biology, University of California, Davis, Davis, CA

**Keywords:** scRNA-seq, data integration, data harmonization, alignment, deep learning, neural networks, response to stimulus, batch effects, domain adaptation

## Abstract

scRNA-seq dataset integration occurs in different contexts, such as the identification of cell type-specific differences in gene expression across conditions or species, or batch effect correction. We present scAlign, an unsupervised deep learning method for data integration that can incorporate partial, overlapping or a complete set of cell labels, and estimate per-cell differences in gene expression across datasets. scAlign performance is state-of-the-art and robust to cross-dataset variation in cell type-specific expression and cell type composition. We demonstrate that scAlign reveals gene expression programs for rare populations of malaria parasites. Our framework is widely applicable to integration challenges in other domains.

## Background

Single cell RNA sequencing (scRNA-seq) technologies enable the capture of high resolution snapshots of gene expression activity in individual cells. As the generation of scRNA-seq data accelerates, integrative analysis of multiple scRNA-seq datasets^1–8^ is becoming increasingly important. The goal of scRNA-seq data integration is to characterize and eliminate the effect of experimental factors driving expression variation between multiple scRNA-seq datasets, so that downstream analyses such as clustering^9,10^ and trajectory inference^10–12^ performed on all datasets jointly are not driven by these factors. Such experimental factors include both technical nuisance factors such as batch or sequencing protocol^13–18^, as well as biological factors of interest such as in case-control studies^19–22^ or speciation^23^.

Integrative analyses are challenging due to several factors. First, dataset integration can be viewed as mapping one dataset onto another. For example, in case-control studies for which a pair of scRNA-seq datasets are generated from biological replicate populations before and after stimulus, functionally matched cell types across datasets must be identified and aligned in order to estimate cell type-specific response to stimulus. The more differential the response of the individual cell types, the more complex a mapping is required. Therefore, integrative tools must be able to freely scale up or down the complexity of their mapping functions to successfully perform integration depending on the heterogeneity of cell type-specific response to stimulus. In the extreme case where some cell types are present in only a subset of conditions being integrated, this poses additional mapping challenges since there may not be a 1-1 correspondence between types across conditions. Second, current integrative tools can be separated into two exclusive sets: those that require all cells from all datasets to have known cell type labels (supervised), and those that do not make use of any cell type labels (unsupervised). Consequently, when only a subset of cells can be labeled with high accuracy, or if only one dataset is labeled (as is the case when reference annotated cell atlases are available^24–29^), this partial set of labels currently cannot be used in data integration. Third, measured transcriptomes even for homogeneous populations of cells occupy a continuum of cell states, for both technical^30,31^ and biological^32–34^ reasons. Thus, individual cells cannot be matched exactly across datasets. Therefore, downstream analysis of integrated datasets typically involves clustering cells across datasets to find matching cell types and estimating cell type-specific differences across datasets. The clustering step makes it difficult to find rare cell populations that differ between datasets.

Here we present scAlign, a deep learning-based method for scRNA-seq alignment. scAlign performs single cell alignment of scRNA-seq data by learning a bidirectional mapping between cells sequenced within individual datasets, and a low-dimensional alignment space in which cells group by function and type, regardless of the dataset in which it was sequenced. This bidirectional map enables users to generate a representation of what the same cell looks like under each individual dataset, and therefore simulate a matched experiment in which the exact same cell is sequenced simultaneously under different conditions. Compared to previous approaches, scAlign can scale in alignment power due to its neural network design, and it can optionally use partial, overlapping, or a complete set of cell type labels in one or more of the input datasets. We demonstrate that scAlign outperforms existing alignment methods including Seurat^3,35^, scVI^7^, MNN^36^, scanorama^8^, scmap^37^, MINT^1^ and scMerge^4^, particularly when individual cell types exhibit strong dataset-specific signatures such as heterogeneous responses to stimulus. While misalignment of cell types unique to one dataset is an inherent challenge for any alignment technique, we show that scAlign produces minimal false positive matchings. Furthermore, we show that our bidirectional map enables identification of changes in rare cell types that cannot be identified from alignment and data analysis steps performed in isolation. We also demonstrate the utility of scAlign in identifying changes in expression associated with sexual commitment in malaria parasites and posit that scAlign may be used to perform alignment in domains other than single cell genomics as well.

## Results

The overall framework of scAlign is illustrated in **Figure 1**. While this paper is written in the context of integrating multiple datasets representing cell populations exposed to different stimuli or control conditions, scAlign can be readily used for any data integration context discussed in the introduction. The premise of integration methods is that when similar cell populations are sequenced under different conditions, some (possibly large) separation can be observed between cells of the same functional type but sequenced in different conditions (**Fig. 1a**). The first component of scAlign is the construction of an alignment space using scRNA-seq data from all conditions, in which cells of the same functional type are indistinguishable, regardless of which condition they were sequenced in (**Fig. 1b**). This alignment space represents an unsupervised dimensionality reduction of scRNA-seq data from genome-wide expression measurements to a low dimensional manifold, using a shared deep encoder neural network trained across all conditions. Unlike autoencoders, which share a similar architecture to scAlign but use a different objective function, our low dimensional manifold is learned by training the neural network to simultaneously encourage overlap of cells in the state space from across conditions (thus performing alignment), yet also preserving the pairwise cell-cell similarity within each condition (and therefore minimizing distortion of gene expression). Optionally, scAlign can take as input a partial or full set of cell annotations in one or more conditions, which will encourage the alignment to cluster cells of the same type in alignment space.

**Figure 1.**
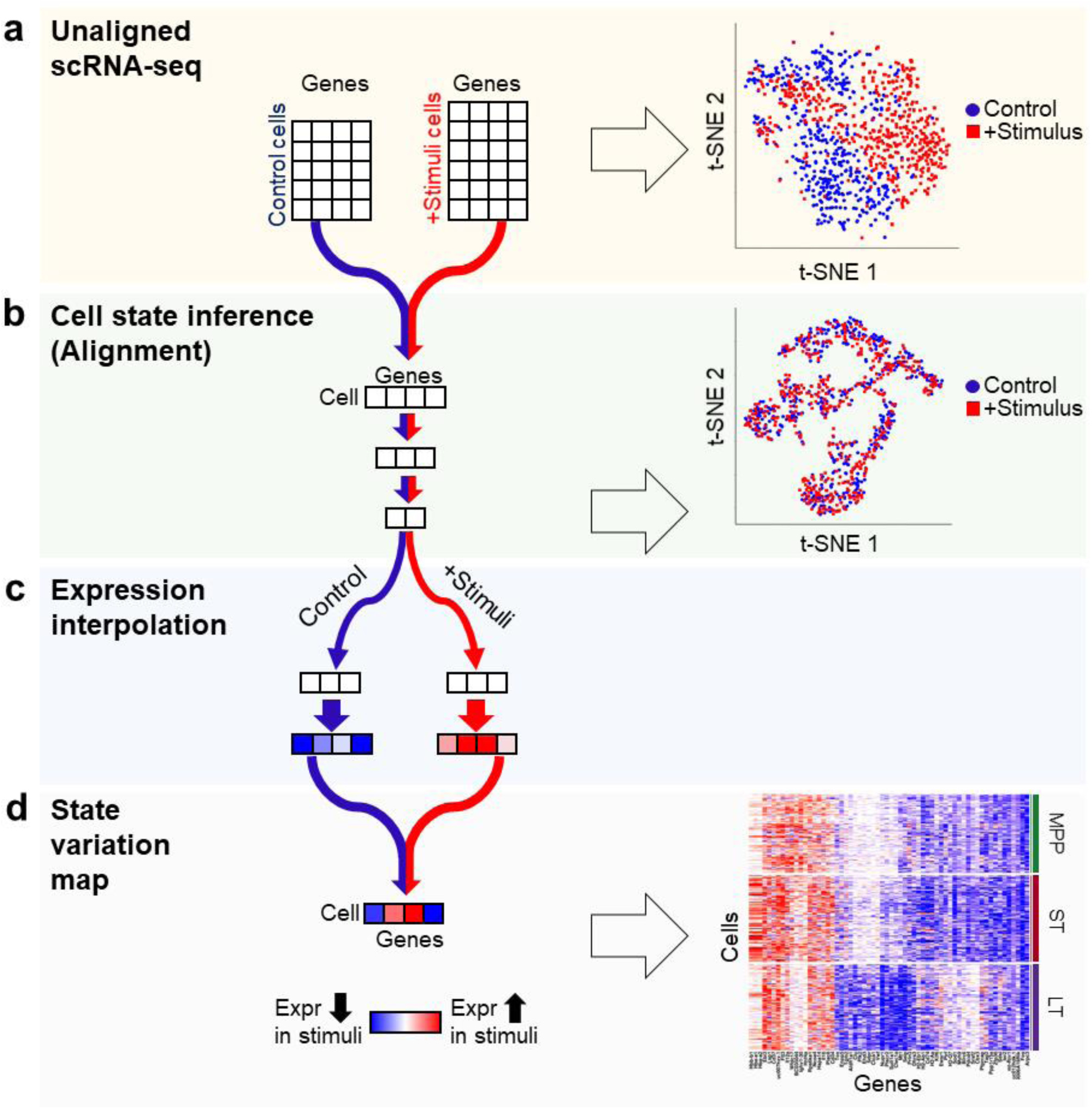
Schematic of unsupervised alignment and state variation mapping with scAlign. (**a**) The input to scAlign consists of cells sequenced across multiple scRNA-seq conditions. Expression can be represented as either gene-level expression, or embedding coordinates from dimensionality reduction techniques such as PCA or CCA. (**b**) A deep encoding network learns a low-dimensional alignment space that simultaneously aligns cells from all conditions. (**c**) Paired decoders project cells from the alignment space back into the gene expression space of each condition, and can be used to interpolate the expression profile of cells sequenced from any condition into any other condition. (**d**) For a single cell sequenced under any condition, we can calculate its interpolated expression profile in all conditions, then measure the predicted variance across all input conditions to calculate a state variation map for the same cell state under different conditions to identify cells whose expression profiles vary significantly across condition.

In the second component of scAlign (**Fig. 1c**), we train condition-specific deep decoder networks capable of projecting individual cells from the alignment space back to the gene expression space of each input condition, regardless of what condition the cell is originally sequenced in. We use these decoders to measure per-cell and per-gene variation of expression across conditions, which we term the cell state variation map. In the case of integrating two conditions, this cell state variation map estimates a paired difference in expression of the same cell across conditions (**Fig. 1d**). scAlign therefore seeks to re-create the ideal experiment in which the exact same cell is sequenced before and after a stimulus in a case-control study, for example.

### Results - scAlign captures cell type specific response to stimulus

We first benchmarked the alignment component of scAlign using data from four publicly available scRNA-seq studies for which the same cell populations were sequenced under different conditions, and for which the cell type labels were obtained experimentally (**Fig. 2, Additional file 1: Fig. S1**). Our first benchmark is CellBench^38^, a dataset consisting of three human lung adenocarcinoma cell lines (HCC827, H1975, H2228) that were sequenced using three different protocols (CEL-Seq2, 10x Chromium, Drop-Seq Dolomite) as well as at varying relative concentrations of either RNA content or numbers of cells in a mixture. While the alignment of the homogeneous cell populations sequenced across protocols was trivial and did not require data integration methods (**Additional file 1: Fig. S2**), alignment of RNA mixtures across protocols was more challenging and more clearly illustrated the performance advantage of scAlign (**Fig. 2a**). We additionally benchmarked alignment methods using data generated by Kowalczyk et al.^39^ and Mann et al.^40^ on three hematopoietic cell types (LT-HSC, ST-HSC, MPP) collected from the C57BL/6 mouse strain at approximately 2 months (“young”) and 2 years (“old”) of age. Mann et al. additionally challenged the mice with an LPS or a control stimulus. Similar to our results with CellBench, scAlign outperforms other approaches on both of these benchmarks (**Fig. 2b**,**c**). The results of scAlign in these comparisons were robust to network depth, width and input features (**Additional file 1: Fig. S3, S4**) along with choice of hyper parameters.

**Figure 2.**
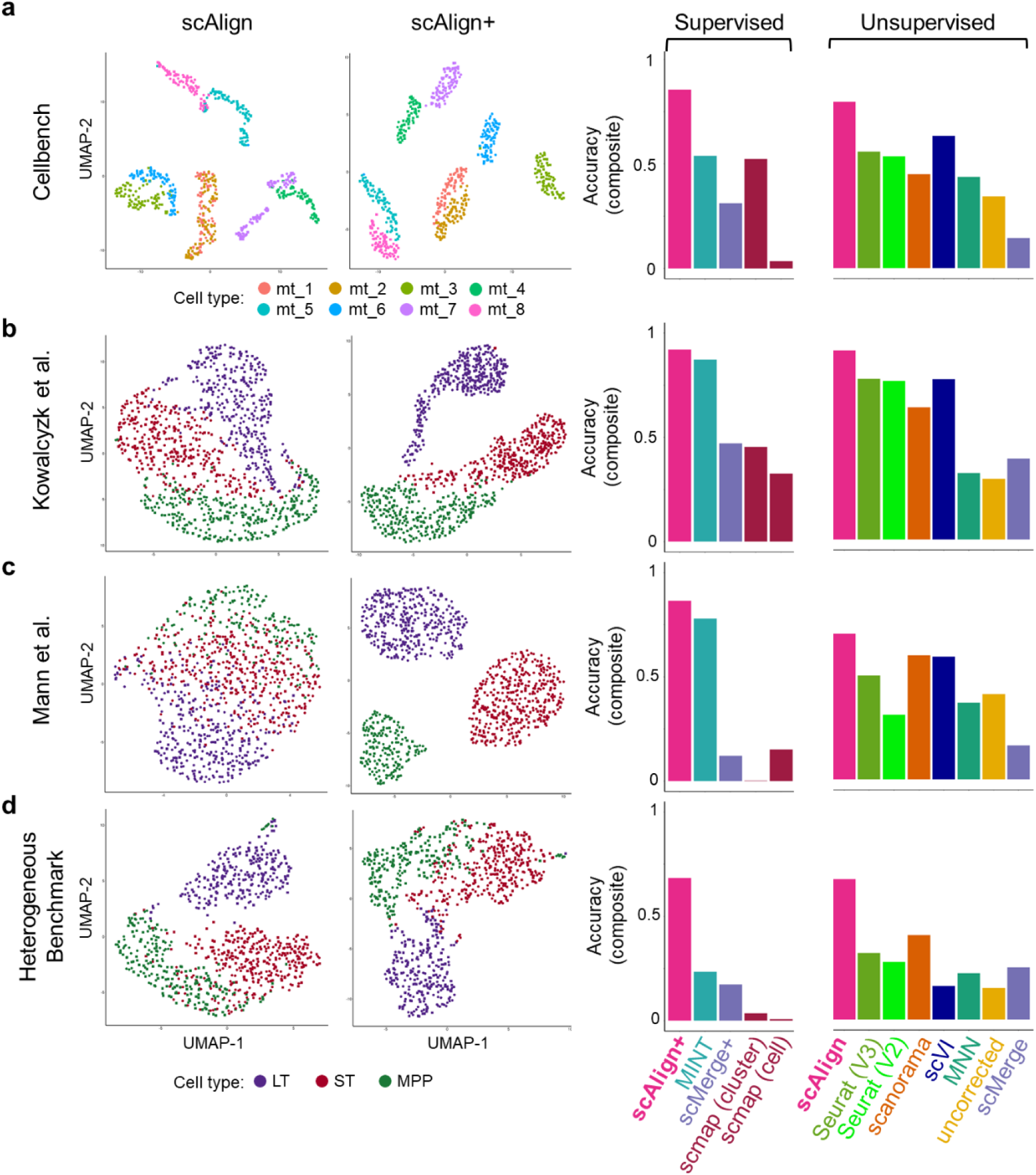
scAlign outperforms existing alignment approaches on four benchmarks. (**a**) CellBench, a benchmark consisting of mixtures (mt) of RNA from three cancer cell lines sequenced using multiple protocols. Plots from left to right: (1) UMAP plot of embeddings after alignment with scAlign, where each point represents a cell, and cells are colored according to their mixture type (mt) as reported in Tian et al. (2) UMAP plot of embeddings after alignment with supervised scAlign (scAlign+). (3) Bar plot indicating the accuracy_composite_ (see Methods) of a classifier, measured as a weighted combination of cross-condition label prediction accuracy and alignment score. (**b**) Same as (a), but with the Kowalczyk et al. benchmark consisting of hematopoietic cells sequenced from young and old mice. Cells are colored according to type (LT, ST, MPP, legend at bottom). (**c**) Same as (a), but with the Mann et al. benchmark consisting of hematopoietic cells sequenced from young and old mice, challenged with LPS. (**d**) Same as (a), but with the HeterogeneousBenchmark dataset consisting of hematopoietic cells responding to different stimuli.

To better understand why the relative performance of the other methods was inconsistent across benchmarks (**Fig. 2a-c**), we next characterized the difficulty of each benchmark for alignment. For each cell type in each benchmark, we identified cell type marker genes by computing the differentially expressed genes (DEGs) between cell types, individually for each condition. We observed considerable overlap in the cell type marker genes (**Additional file 1: Fig. S5**), suggesting these benchmarks may be less challenging to align and therefore more difficult to distinguish alignment methods from each other. We therefore constructed a novel benchmark termed HeterogeneousBenchmark by combining published scRNA-seq data on hematopoietic cells measured across different studies and stimuli. This benchmark yields smaller overlap in cell type marker genes (**Additional file 1: Fig. S5**), which makes it more challenging to align. On HeterogeneousBenchmark, we find that scAlign’s performance is robustly superior, while Seurat and Scanorama also outperform the remaining methods (**Fig. 2d**).

scAlign simultaneously aligns scRNA-seq from multiple conditions and performs a non-linear dimensionality reduction on the transcriptomes. This is advantageous because dimensionality reduction is a first step to a number of downstream tasks, such as clustering into putative cell types^41^ and trajectory inference^42–44^. Dimensionality reduction of cell types generally improves when more data is used to compute the embedding dimensions, and so we hypothesized that established cell types will cluster better in scAlign’s embedding space in part due to the fact we are defining a single embedding space using data from multiple conditions. We therefore compared the clustering of known cell types in the scAlign embedding space to an autoencoder neural network that uses the same architecture and number of parameters as scAlign, but is trained on each condition separately (see Methods). In two of the three benchmarks we tested, we found that known cell types cluster more closely and are more distinct in scAlign embedding space compared to that of the corresponding autoencoder (**Fig. 3, Additional file 1: Fig. S6**), suggesting scAlign’s embedding space benefits from pooling cells from across all conditions. Furthermore, by pooling cells into a common embedding space scAlign can identify new subpopulations within known cell type clusters (**Additional file 1: Fig. S7**).

**Figure 3.**
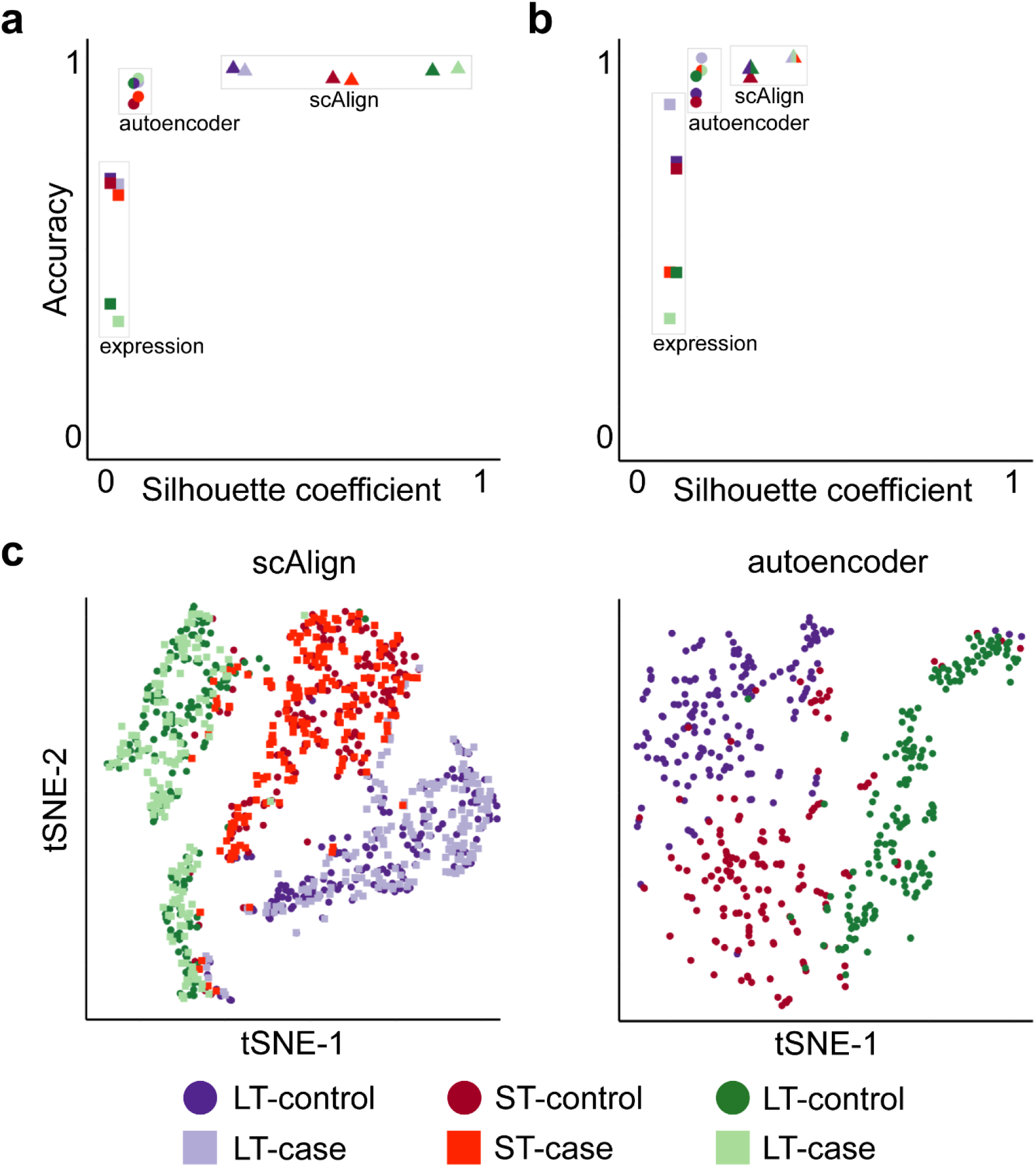
Joint analysis of cells from all conditions leads to more accurate clustering of cell types compared to independent analysis of individual conditions. (**a**) Scatterplot illustrating the quality of clustering of cell types within each condition from the Mann et al. benchmark. Each point represents one cell type in one condition, when the embedding is computed using either the original expression data (‘expression’), the embedding dimensions of scAlign, or the embedding dimensions of an autoencoder with the same neural network architecture as scAlign. The y-axis represents classification accuracy, while the x-axis represents the silhouette coefficient. (**b**) Same as (a), but for HeterogeneousBenchmark (**c**) tSNE plots visualizing the embedding space of scAlign trained on both conditions and (**d**) an autoencoder trained on a single condition.

A unique feature of scAlign is that it can optionally use cell type labels for a subset of (or all) cells if available, but does not require any labels by default. In other words, scAlign can perform unsupervised, semi-supervised or fully-supervised alignment. One example of a use case would be when a labeled, highly quality cell atlas is available, it can be used to label cells sequenced from a newer, smaller study. **Figures 2a-d** illustrate, for each of the four benchmarks, that scAlign performance improves when cell type labels are available at training time, and exceeds the performance of other supervised methods such as MINT^31^, scMerge^4^ and scmap^37^. Even when only a subset of cells from one condition have labels available for semi-supervised training, scAlign performance improves compared to a strictly unsupervised alignment, though still lower than a fully supervised scAlign+ (**Fig. 4, Additional file 1: Fig. S8**). When provided with labels, the cell-cell similarity matrix of the supervised scAlign method is qualitatively similar to the cell-cell similarity matrix of cells in the original gene expression space as well as the unsupervised scAlign alignment space, suggesting the inferred alignment space is robust to adding labels during alignment (**Additional file 1: Fig. S9**).

**Figure 4:**
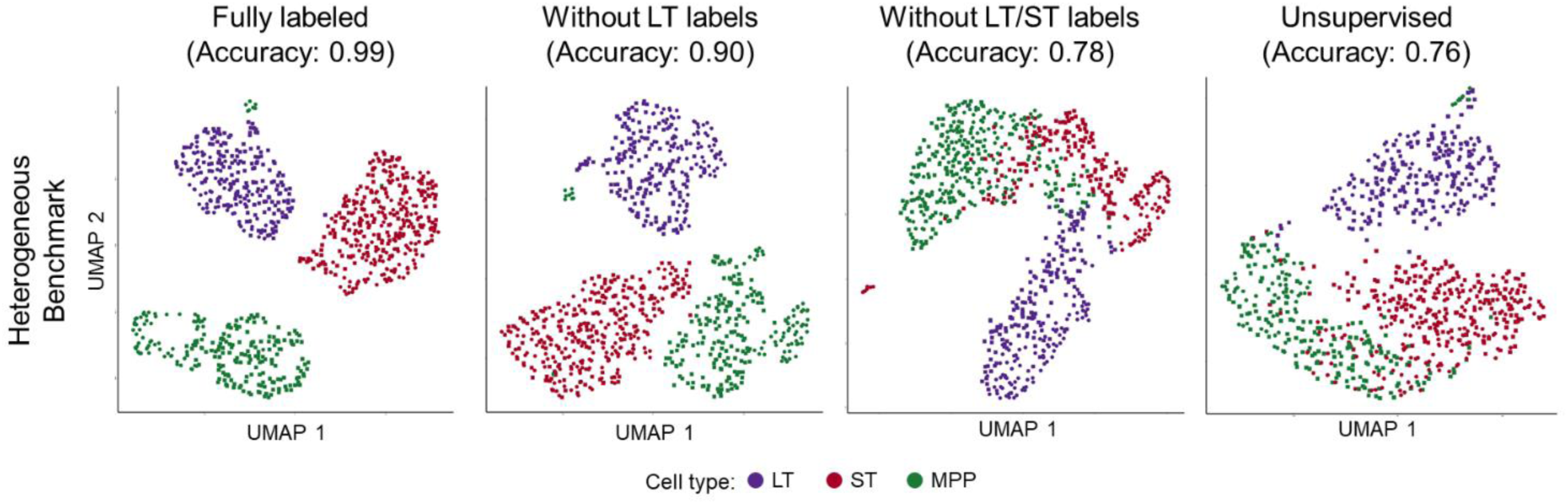
Semi-supervised alignment mode of scAlign enables use of partial sets of cell type labels. UMAP visualization of the HeterogenousBenchmark after alignment with scAlign+ trained with (**a**) labels for all cells in both conditions, (**b**) after removal of labels for LT-HSC HSC in the stimulated condition, (**c**) after removal of labels for LT-HSCs and ST-HSCs in the stimulated condition, and (**d**) scAlign trained without cell labels.

### Results - scAlign is robust to large differences in cell type representation across conditions

Besides cell type-specific responses to stimuli, we reasoned that the other factor that determines alignment difficulty is the difference in the representation (or proportion present) of each cell type across conditions. For example, cell types unique to one condition may pose challenges to alignment because there are no functionally matched cell types in the other conditions. We therefore explored the behavior of scAlign and other approaches when the relative proportion of cell types varies significantly between the conditions being aligned.

We performed a series of experiments on the Kowalczyk et al. benchmark where we measured alignment performance of all methods as we removed an increasing proportion of cells from each cell type from the old mouse condition (**Fig. 5**). While scAlign had superior performance across all experiments and was most robust to varying cell type proportions, surprisingly, we found that other methods were generally robust as well. Removing even 75% of the cells of a given type only led to a median drop of 11% in accuracy across the tested methods. When we repeated these experiments on the Mann et al. benchmark, we generally found a larger decrease in performance as we removed more cells from each type compared to the Kowalczyk et al. benchmark, though scAlign still outperformed all other methods (**Additional file 1: Fig. S10**).

**Figure 5.**
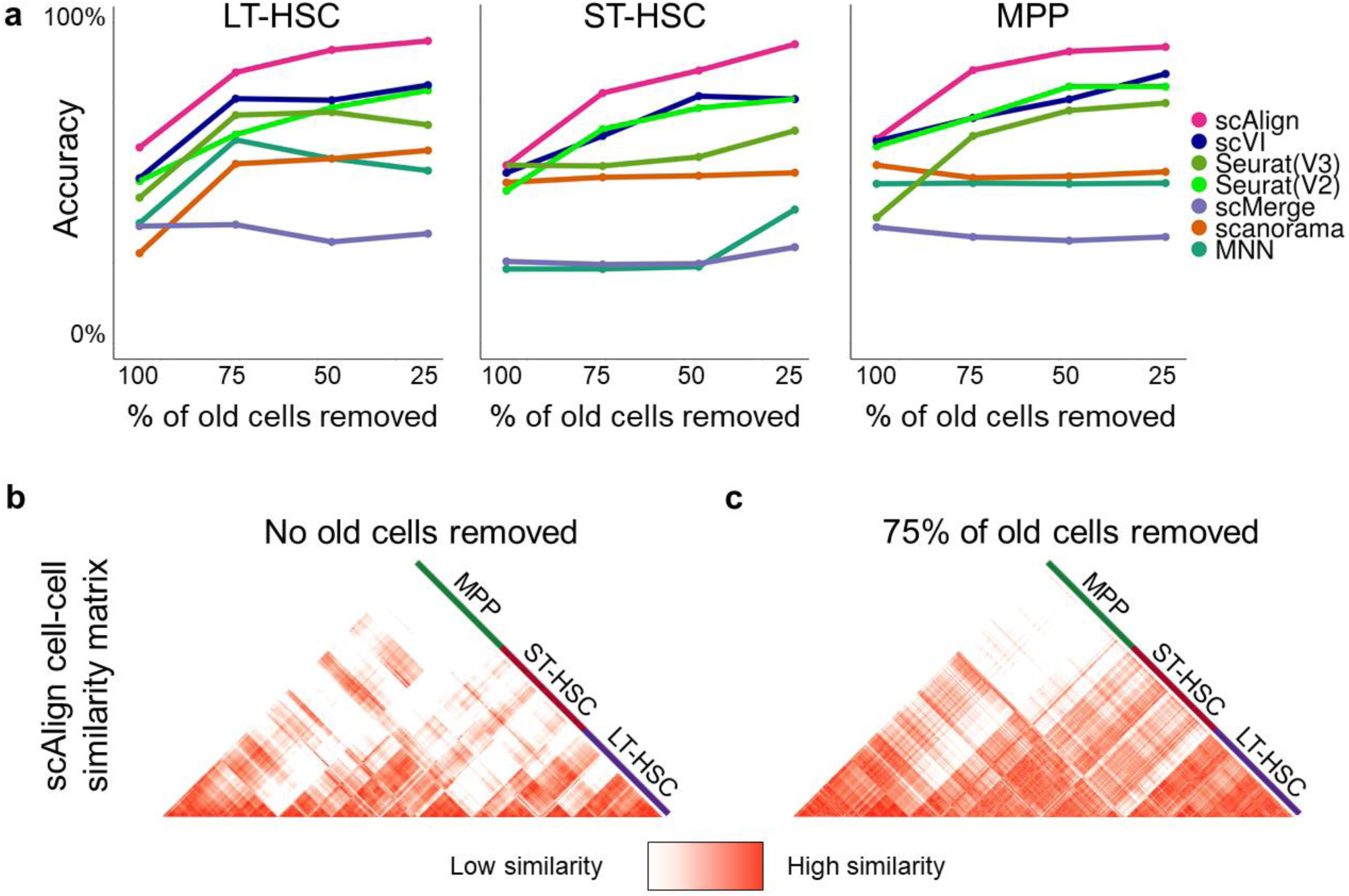
Alignment performance is robust to imbalance in cell type representation across conditions. (**a**) Accuracy of classifiers on the Kowalczyk et al. benchmark, when removing either LT-HSC, ST-HSC or MPP cells from the old condition. scAlign outperforms all other methods and exhibits minimal degradation in performance as increasing numbers of cells are removed within each cell type. (**b**) Heatmap showing the pairwise similarity matrix for the young cells from Kowalczyk et al. when no cells have been removed. (**c**) Heatmap showing the pairwise similarity matrix for the young cells from Kowalczyk et al. after removing 25% of the old mouse cells from all cell types.

We next investigated the factors that underlie scAlign’s robustness to imbalanced cell type representation across conditions. scAlign optimizes an objective function that minimizes the difference between the pairwise cell-cell similarity matrix in gene expression space, and the pairwise cell-cell similarity matrix implied in the alignment space when performing random walks of length two (**Fig. 6a**). The random walk starts with a cell sequenced in one condition, then moves to a cell sequenced in the other condition based on proximity in alignment space. The walk then returns to a different cell (excluding the starting cell) in the original condition, also based on proximity in alignment space. For every cell in each condition, we calculated the frequency that such random walks (initiated from the other condition) pass through it (**Fig. 6b-c**). We found that a select few representatives for each cell type are visited much more frequently than others, and that even when those cells are removed from the condition, another cell is automatically selected as a replacement (**Additional file 1: Fig. S11**). This suggests that a given cell type in one condition only depends on a few cells of the same type in the other condition to align properly, and so scAlign alignment does not need every cell type to be represented in the same proportion across conditions.

**Figure 6.**
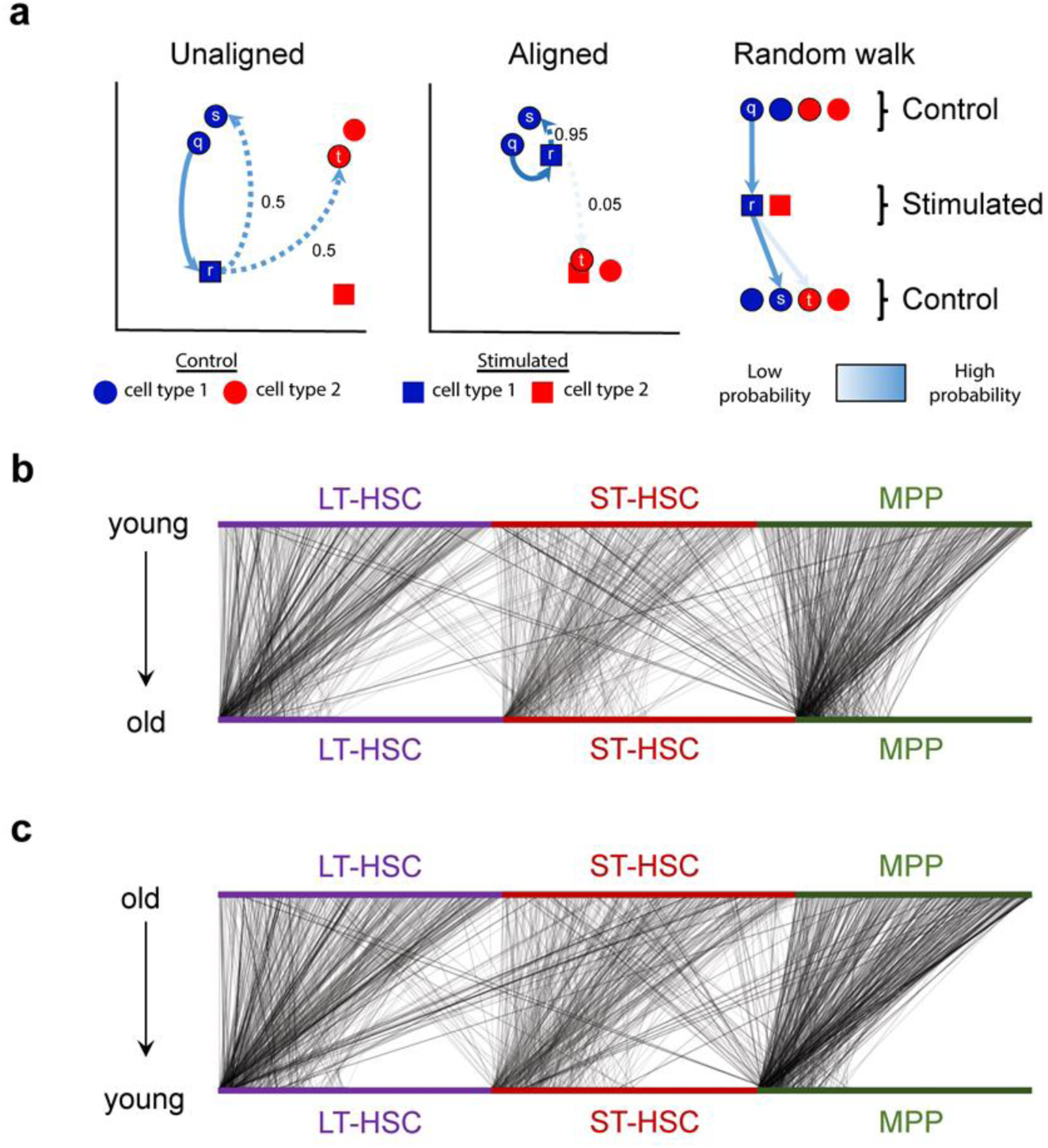
Random walks during scAlign training frequently visit a small number of hub cells. (**a**) Schematic of the cross condition round trip random walk prior to and after training of scAlign. (**b**) Visualization of the probability of a walk from each individual young cell (top) to each individual old cell (bottom) after training scAlign on the Kowalczyk et al. benchmark. Edge density represents the magnitude of the probability of a given walk. (**b**) Same as (a), except the edges represent the probability of walking from individual old cells (top) to individual young cells (bottom) in the Kowalczyk et al. benchmark.

In the above experiments, we have aligned conditions in which the same set of cell types are present in all conditions. We next explored the behavior of scAlign and other approaches when there are cell types represented in only a subset of the conditions. We expect such scenarios to arise when only a subset of cell types respond to, or are targeted by, a stimulus or condition. For each of our benchmarks, we removed one cell type from one of the conditions (e.g. the LPS condition of the Mann benchmark, or the old mouse condition of the Kowalczyk benchmark), and aligned the control and stimulated conditions to determine the extent to which the unique population maintained separation from other cell types after alignment. **Figure 7a** demonstrates that in eight out of nine cases, scAlign outperforms other alignment methods in terms of classification accuracy. Even in cases where the alignment accuracy was similar between methods, scAlign visually separates cell types in its alignment space more so than other approaches such as Scanorama and Seurat (**Fig. 7b**). For other approaches, the separation of different cell types within the same condition shrinks when one cell type is removed (**Additional file 1: Fig. S12**).

**Figure 7.**
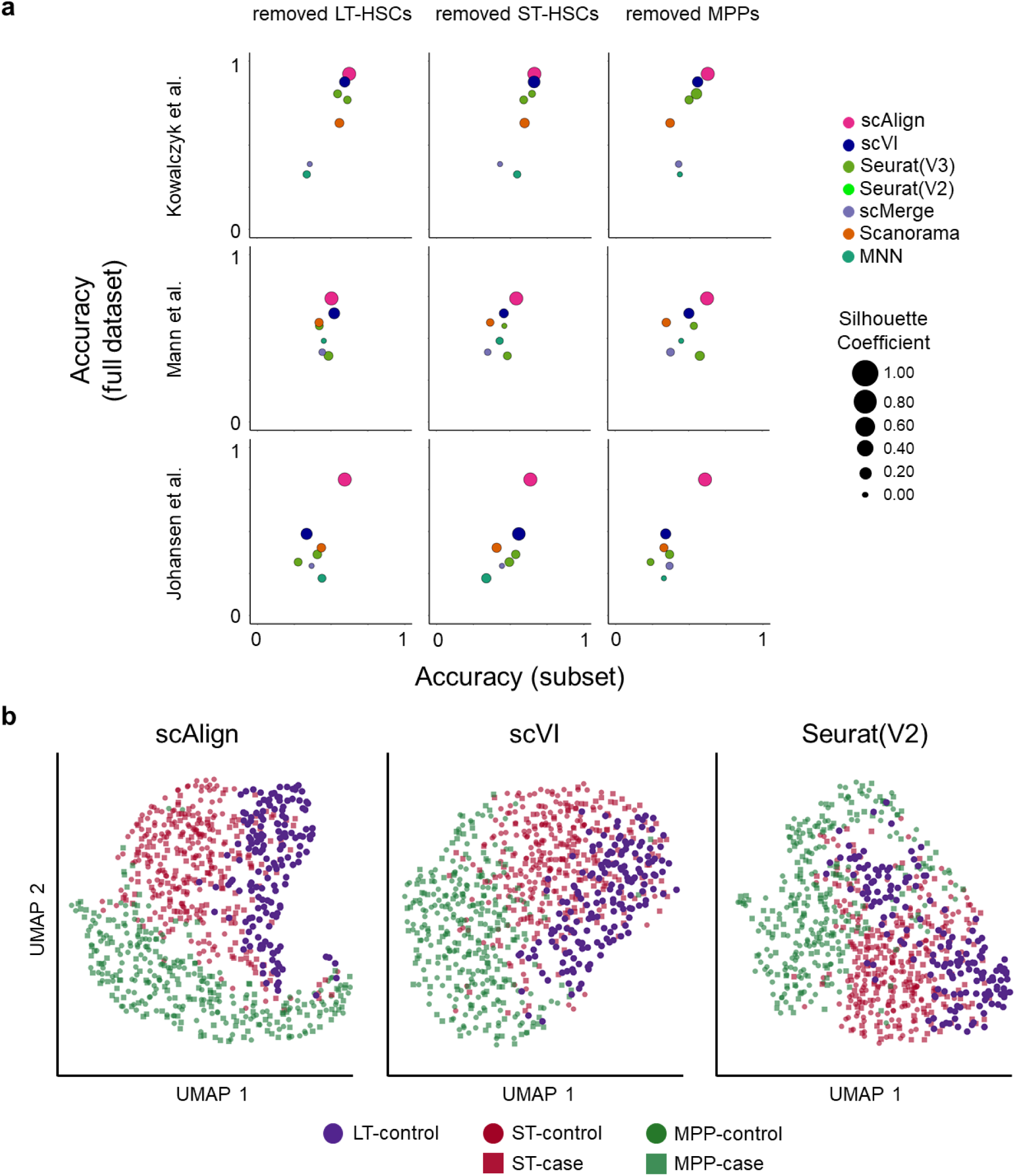
scAlign is robust to distinct cell type sets between conditions. (**a**) Scatterplot matrix of performance of each method when both conditions have the same number of cell types (y-axis), compared to when one cell type has been removed (the LPS condition of the Mann benchmark, or the old mouse condition of the Kowalczyk benchmark) (x-axis). Each point is scaled in size by the silhouette coefficient for the clustering after alignment. (**b**) tSNE plots with cells colored by cell type and condition for the top performing methods.

### Results - scAlign interpolates gene expression accurately

One of the more novel features of scAlign is the ability to map each cell from the alignment space back into the gene expression space of each of the original conditions, regardless of which condition the cell was originally sequenced in. This mapping is performed through interpolation: for each condition, we learn a mapping from the alignment space back to gene expression space using cells sequenced in that condition, then apply the map to all cells sequenced in all other conditions. This interpolation procedure enables measurement of variation in gene expression for the same cell state across multiple conditions, and simulates the ideal experiment in which the exact same cell is sequenced before and after a stimulus is applied, and the variation in gene expression is subsequently measured.

To measure the accuracy of scAlign interpolation, for each of the three hematopoietic benchmarks, we trained decoder neural networks to map cells from the alignment space back into each of the case and control conditions. We then measured interpolation accuracy as the accuracy of a classifier trained on the original gene expression profiles of cells sequenced under one condition (e.g. stimulated), when used to classify cells that have been interpolated from the other condition (e.g. control). Comparing this interpolation accuracy to cross-validation accuracy of classifying cells in their original condition using the original measured gene expression profiles, we see that interpolation accuracy is similar to expression accuracy (**Fig. 8a**), suggesting that cells maintain their general type when mapped into another condition.

**Figure 8.**
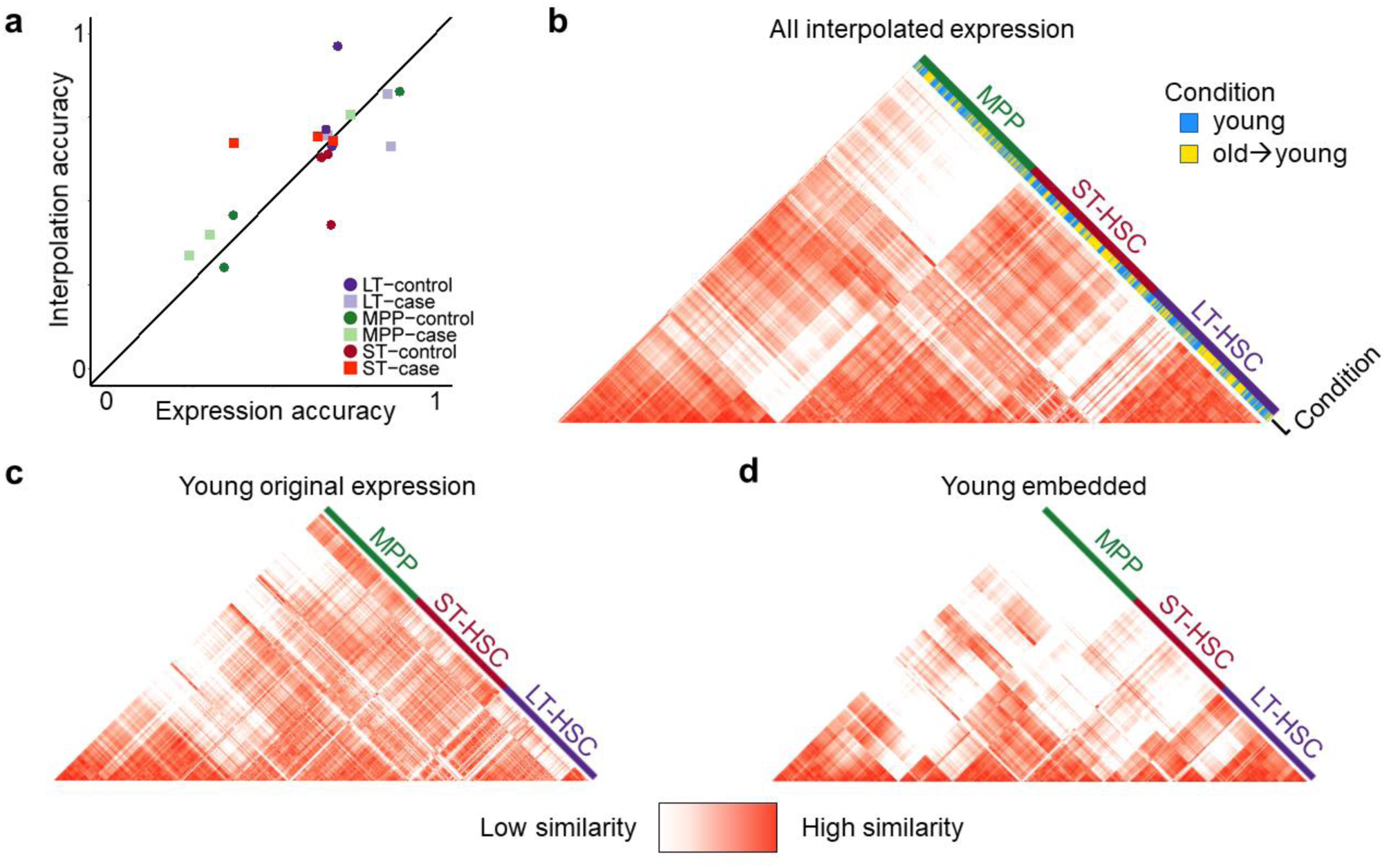
Interpolation of gene expression patterns is accurate. (**a**) Scatterplot of classifiers trained on gene expression profiles of one condition, that are subsequently used to predict labels of either measured expression profiles from the same condition in a cross-validation framework (x-axis), or used to predict labels of cells sequenced from the other condition that were then interpolated into this condition (y-axis). Similarity in accuracy represented by points near the diagonal indicates that cell type identity encoded in the gene expression profile is maintained even after interpolation. (**b**) The pairwise cell-cell similarity matrix for all cells projected into the young condition, including both the old cells interpolated into the young condition (yellow) and the cells originally sequenced in the young condition (blue). Note that cells cluster largely by cell type regardless of the condition in which they were sequenced. (**c**) The pairwise cell-cell similarity matrix for all cells computed using the original expression measurements. (**d**) The pairwise cell-cell similarity matrix for all cells computed using the low-dimensional coordinates within the alignment space learned by scAlign. Similarity between (c) and (d) indicate the scAlign embedding maintains global similarity patterns between cells in the original gene expression space.

**Figure 8b** illustrates the cell-cell similarity matrix computed in gene expression space of hematopoietic cells collected in the Kowalczyk study, when including cells sequenced in the young mice, as well as cells that have been interpolated from the old mice into the young condition. We see that cells cluster largely by cell type (LT-HSC, ST-HSC, MPP) and not by their condition of origin. Furthermore, by computing a state variance map from the interpolation of all cells into both conditions, we identify differentially expressed genes that were not identified by traditional differential expression analysis (**Additional file 1: Fig. S13**). This demonstrates that the encoding and interpolation process maintains data fidelity, even though the encoder is trained to align data from multiple conditions and is not explicitly trained to minimize reconstruction error like typical autoencoders. **Figures 8c**,**d** further illustrate that the cell-cell similarity matrix in embedding space is faithful to the cell-cell similarity matrix in the original gene expression space.

### Results – interpolation identifies early gametocyte markers of the engineered ap2-g-dd strain of P. falciparum

We next applied scAlign to identify genes associated with early steps of sexual differentiation in *Plasmodium falciparum*, the most widespread and virulent human malaria parasite. Briefly, the clinical symptoms of infection are the result of exponential growth of asexual parasites within red blood cells, while parasite transmission depends on the formation of the non-replicating male and female sexual stages necessary for infection of the parasite’s mosquito vector. During each round of asexual replication, a sub-population of parasites will activate expression of the *ap2-g* gene, which encodes the transcriptional master regulator of sexual differentiation, to initiate sexual differentiation. While the gene *ap2-g* is a known master regulator of sexual commitment, and its expression is necessary for sexual commitment, the events which follow *ap2-g* activation and lead to full sexual commitment are unknown^45^. Furthermore, *ap2-g* expression is restricted to a minor subset of parasites, making the identification of the precise stage of the life cycle when sexual commitment occurs a challenging task.

**Figure 9a** illustrates the alignment space of parasites which are either capable of *ap2-g* expression and will contain an *ap2-g*-expressing subpopulation in the initial stages of sexual differentiation (+Shld), or are *ap2-g* deficient and therefore all committed to continued asexual growth (−Shld). As was observed in the original paper^45^, the +/−Shld cells fall into clusters that can be ordered by time points in their life cycle (**Fig. 9a**). scAlign alignment maintains the gametocytes from the +Shld condition as a distinct population that is not aligned to any parasite population from the −Shld condition, whereas other tested methods are unable to isolate the gametocyte population (**Additional file 1: Fig. S14**).

**Figure 9.**
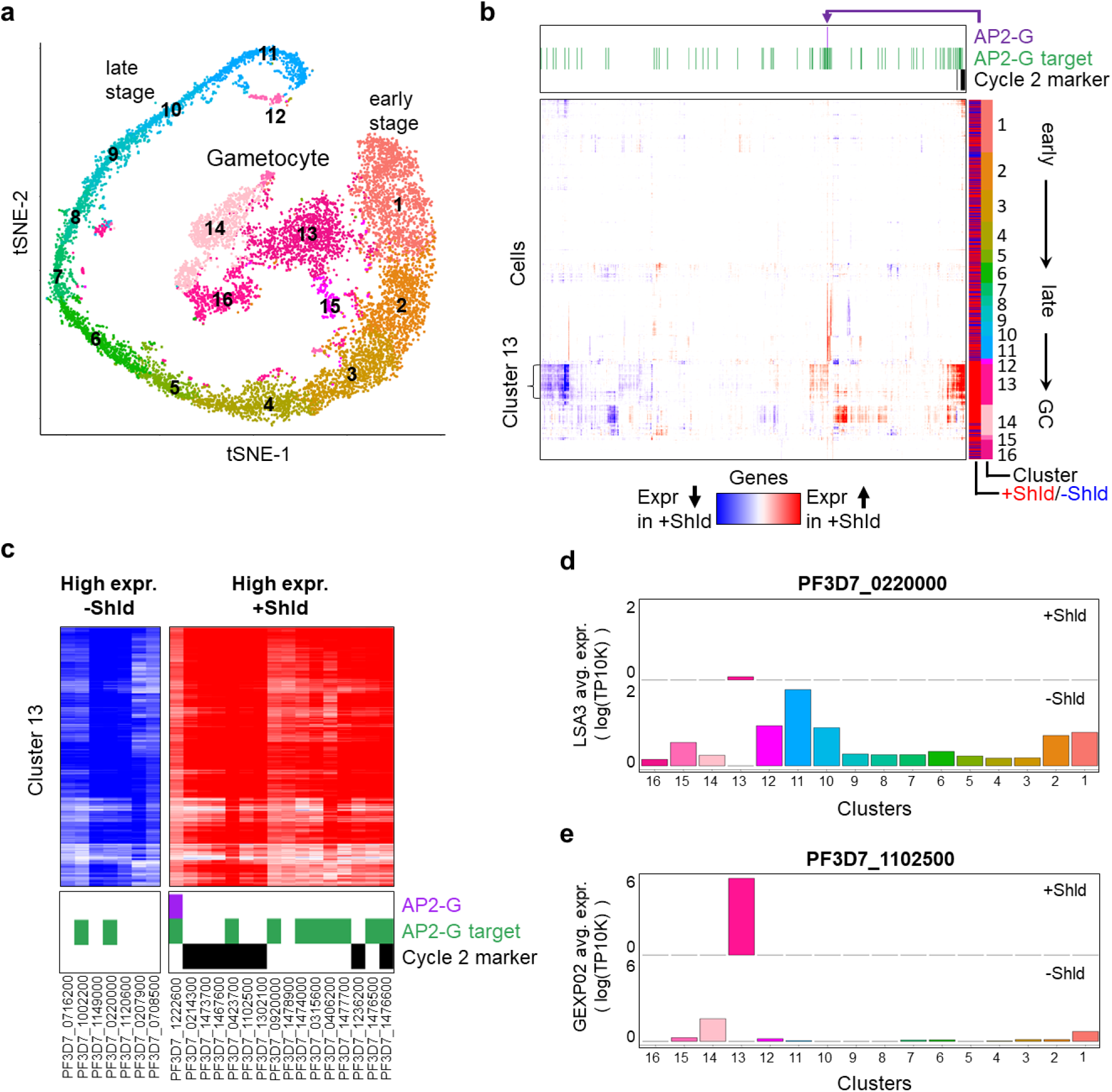
Alignment of *P. falciparum* cells sequenced from a conditional ap2-g knockdown line identifies cycle 2 gametocytes. (**a**) tSNE visualization of cells that cannot stably express *ap2-g* (−Shld) and *ap2-g* expression-capable cells (+Shld) after alignment by scAlign. Each cell is colored by its corresponding cluster identified in Poran et al., and clusters are numbered according to relative position in the parasite life cycle. (**b**) scAlign state variation map defined by projecting every cell from (a) into both the +/−Shld conditions, then taking the paired difference in interpolated expression profiles. Rows represent cells, ordered by cluster from early stage (top) to late stage and GC (bottom), and columns represent the 661 most varying genes. The state variation map reveals that cluster 13 is predicted to differ in expression the most between +/−Shld. The column annotations on top indicate which of the variable genes have been previously established as a target of *ap2-g* via ChIP-seq experiments^47^ which genes have been reported as playing a role in cell cycle 2 gametocyte maturation^46^ and which gene represents *ap2-g*. (**c**) The same state variation map of (c), but zoomed in on Cluster 13 and the genes predicted to be most differentially expressed between +/−Shld. (**d**) Average per-cluster expression levels of PF3D7_0220000 reported in (c), for both the +/−Shld conditions. PF3D7_0220000 is predicted to be up-regulated in −Shld relative to +Shld, which is reflected in the per-cluster expression levels. (**e**) Same as (d), but for PF3D7_1102500, a gene predicted to be up-regulated in +Shld relative to −Shld.

To further investigate how scAlign is able to maintain the gametocytes as a distinct population after alignment, we looked at the random walks performed by the gametocyte cells to see which cells from the −Shld condition they walked to, and found that scAlign maps a very small number of cells from similar surrounding clusters into the peripheral region of alignment space near the gametocytes. These −Shld cells in the periphery of the gametocyte cluster allows the gametocytes to use those cells as “anchors” in their random walk and maintain their overall separation from the −Shld cells. To confirm this hypothesis, we removed the contaminating −Shld parasites used as anchors by the +Shld gametocytes, and re-aligned the +Shld and reduced set of −Shld cells. After realignment, we found that scAlign “sacrificed” parasites from similar surrounding clusters to act as new anchors and preserve the distinct +Shld gametocytes as a distinct population (**Additional file 1: Fig. S15**).

Because the +Shld and −Shld cells form a set of clusters that we could order from early stage to late stage then gametocytes (+Shld), we hypothesized that the state variation map computed by scAlign could reveal where in the life cycle sexual-committing cells (a subset of +Shld cells) distinguished themselves in variation from asexual-committing cells (all −Shld cells). Using the interpolation component of scAlign, we projected each cell sequenced from each condition in the alignment space into the expression space of both of the +/−Shld conditions. By taking the difference in interpolated expression for each cell between the +Shld and −Shld transcriptomes, we computed a state variation map illustrating the predicted difference between the two conditions along the entire life cycle (**Fig. 9b**). From the state variation map, we observed few overall predicted differences in gene expression between the two conditions across most stages of the life cycle, except within a cluster of cells containing the gametocytes specific to the +Shld condition (**Fig. 9b**, cluster 13). In other words, gametocytes from cluster 13 exhibited the largest predicted differential gene expression between the +Shld gametocytes and neighboring −Shld non-gametocyte parasites. We verified that scAlign interpolation uses cells from neighboring clusters to predict −Shld expression within cluster 13 (**Fig. 9d**,**e**, see Methods).

Over all 661 highly variable genes we analyzed, we found the predicted differentially expressed genes in cluster 13 are enriched in genes previously established to play a role in gametocyte maturation (**Fig. 9b**) (*p* = 1.2 × 10^−6^, Wilcox rank sum test), including *pfg27* (PF3D7_1302100) and *etramp4* (PF3D7_0423700)^46^. Furthermore, for the genes we predict to be upregulated in cluster 13 of the +Shld condition, we observed an enrichment of *ap2-g* targets identified via ChIP-Seq^47^ (*p* = 6.8 × 10^−7^, Wilcox rank sum test). This upregulation of *ap2-g* targets is consistent with the fact that cells that have entered the gametocyte stage must have turned on *ap2-g* expression, but that Shld-cells cannot express *ap2-g*. Our state variation map identifies an additional eight genes not reported by Bancells and colleagues as playing a role in gametocyte maturation, but that are predicted to differ between +/−Shld (**Fig. 9c**). Taken in total, these results suggest the other genes we have predicted as differing between +/−Shld may also play a role in gametocyte conversion (**Fig 9b**,**c**).

### Results – scAlign identifies highly variable genes in pancreatic islet cells sequenced using multiple protocols

We next tested scAlign’s ability to infer an alignment space across more than two conditions by aligning pancreatic islet cells^15^ derived from 8 donors and captured using four different protocols (CEL-Seq, CEL-Seq2, Smart-Seq2 and C1). The un-aligned pancreatic islet cells separate by protocol and not cell type, indicating strong protocol-specific effects which are removed after scAlign alignment (**Additional file 1: Fig. S16, S17a).** scAlign outperforms Seurat and scVI in terms of composite alignment accuracy on this dataset (**Additional file 1: Fig. S17b-c**). Interestingly, scAlign preserves the stellate, ductal and gamma cell types as separate clusters of cells, even though these three groups are represented in only a subset of the four protocols.

Having aligned the pancreatic islet cells into an alignment space, we next computed scAlign’s state variance map to identify cell types and genes exhibiting high expression variation across three protocols to provide insight into how the choice of protocol affects gene expression measurement (**Fig. 10a-d**). Here we excluded C1 because of the overall high gene expression specific to this protocol. We identified multiple subpopulations of cells within the alpha and beta cell types that are remarkably variable across protocols (**Fig. 10e**). We further show that our state variance map identifies subpopulations of alpha cells that are not consistent with the subclustering of alpha cells based on the embeddings (alignment space), illustrating that the state variance map finds unique patterns of expression variation across conditions not found by classic clustering approaches (**Fig. 10f**). Notably, the most highly variable genes with respect to protocol were specific to the activated stellate cells, and we confirmed these genes to be enriched in gene functions related to stellate function (**Additional file 2**).

**Figure 10:**
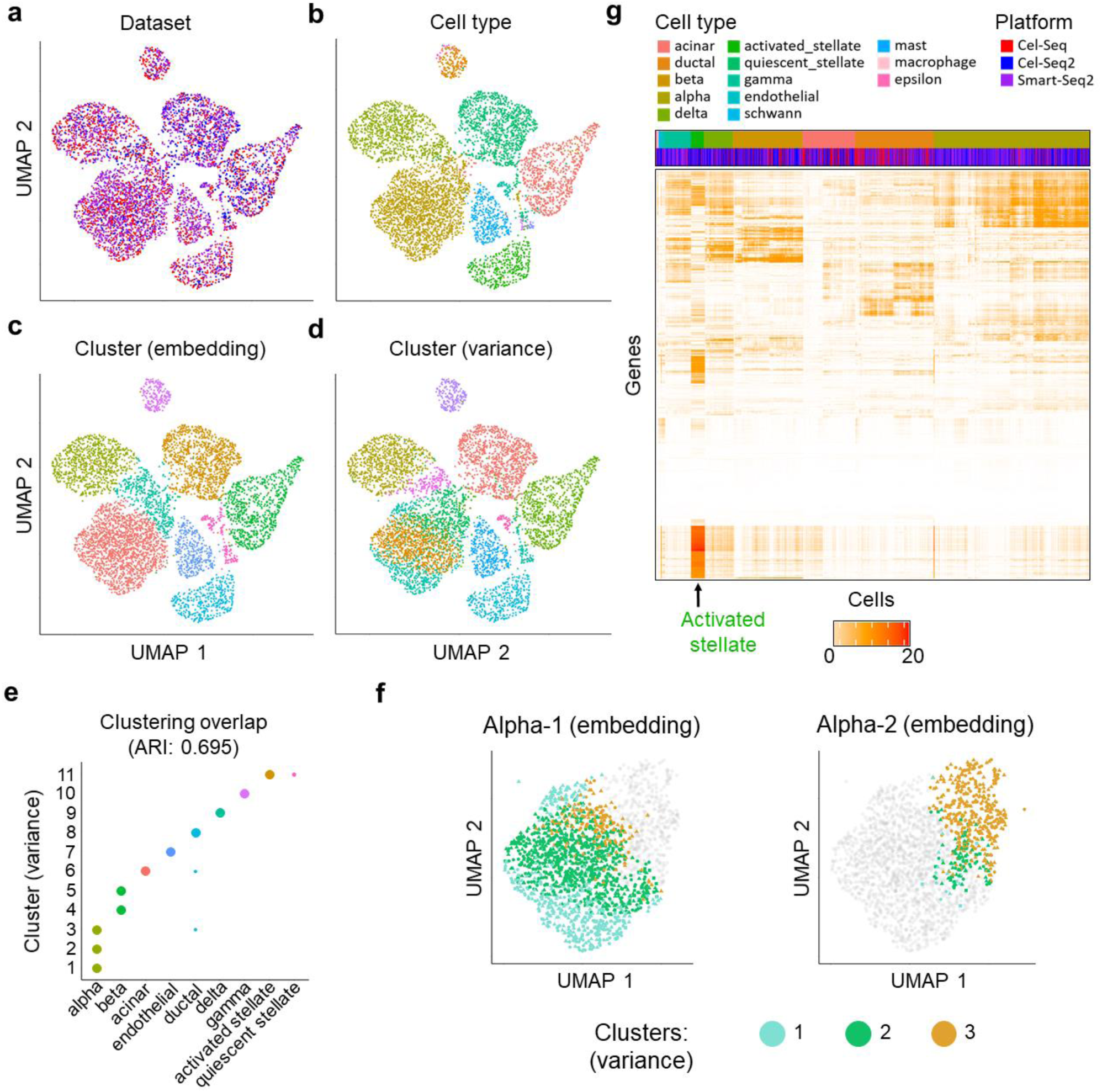
Alignment of pancreatic islet cells captured using three different protocols identifies cell type specific variation across protocols. (**a-d**) UMAP visualization of pancreatic islet cells sequenced on CEL-Seq, CEL-Seq2 and Smart-Seq2 after alignment by scAlign. colored by protocol, cell type, clustering on the alignment space or scAlign’s state variance map. (**e**) Scatterplot indicating the overlap of clusters defined using the state variance map (y-axis) and based on the cell type labels as reported in Stuart et al. (**f**) Comparison of clusters identified using the embeddings, versus using the state variance map. Shown are two clusters defined in the embedding space, termed alpha-1 and alpha-2 because of their overlap with the alpha cell type. Grey points in the alpha-1 plot indicate cluster 2 cells, and grey points in the alpha-2 plot indicate cluster 1 cells. Colored points represent the three clusters identified in the state variance map. scAlign’s variance map clusters (1, 2 and 3) are each found in both alpha-1 and alpha-2, indicating poor agreement. (**g**) Heatmap of the state variance map computed across the three capture protocols (CeL-Seq, CEL-Seq2 and Smart-Seq2) where red indicates high variance of expression predicted for a given gene and cell across protocols

### Results – Robust cell type marker genes drive alignments

To gain insight into the general principles and genes used by scAlign to perform alignment, we performed a series of *in silico* expression perturbation experiments. scAlign uses the same feed-forward network to reduce the dimensionality of cells from all input conditions. We therefore hypothesized that scAlign is implicitly identifying cell type marker genes that are invariant (robust) across conditions, and using these marker genes to perform dimensionality reduction as they will naturally cause similar cell types across conditions to map to the same regions of alignment space. We tested this hypothesis by first identifying a set of marker genes for each cell type that were robust across conditions within a given dataset (**Additional file 3**) (see Methods). We then systematically perturbed the expression of all common marker genes across all cells, and measured the downstream effect of the perturbation on the embeddings of the cells in alignment space. Intuitively, perturbing the expression levels of genes that more strongly contribute to the alignment will yield larger deviations in the embeddings of the cells. As a control, we performed the same perturbation experiments on random control sets of genes matched for size and expression level (see Methods). Perturbing the common marker genes yielded significantly larger deviations in the cell embeddings than the control sets (P < 10^−4^, Permutation test), with the embeddings moving an average of 5.2 fold more than the control sets (**Additional file 1: Fig. S18**).

We next sought to evaluate the extent to which scAlign’s unique random walk-based objective function contributes to its alignment accuracy. Traditional neural networks that focus on unsupervised dimensionality reduction such as autoencoders use an objective function that explicitly learn embeddings that minimize the reconstruction loss of each cell. In contrast, the scAlign objective function simultaneously encourages embeddings to maintain cell-cell similarity within condition, as well as match cells in the alignment space across conditions. We therefore evaluated the utility of scAlign’s objective function by substituting scAlign’s loss function for a classic reconstruction loss-based autoencoder loss function. This autoencoder shares the same number of layers and nodes per layer as scAlign, and furthermore uses a shared encoder across all conditions similar to scAlign, but unique decoders for each condition (see Methods). Both the autoencoder and scAlign therefore have the same number of parameters and therefore equal model capacity, and only differ by their respective objective functions. When comparing this autoencoder to scAlign on each of our four benchmarks, we found that the autoencoder was able to achieve similar accuracy on benchmarks with minimal cell type-specific condition effects, such as Cellbench and Kowalczyk et al. (**Additional file 1: Fig. S19a-b**). However, on more challenging benchmarks such as Mann et al. and our HeterogeneousBenchmark, the autoencoder performed worse than scAlign (**Additional file 1: Fig. S19c-d**). Furthermore, the autoencoder did not maintain the cell-cell similarity matrix in embedding space as well as scAlign (**Additional file 1: Fig. S19, S20**), suggesting the low dimensional embeddings learned by the autoencoder may not as faithfully recapitulate the gene expression inputs.

## Discussion

We have shown that scAlign outperforms other integration approaches, particularly when there are strong cell type-specific differences across conditions, or when there is an imbalance in cell type representation across conditions. Compared to other approaches, scAlign will be particularly useful in the context where only some cell type labels are available in one or more conditions. We envision two scenarios where this may occur. First, with the increasing number of cell atlases^24–29^ that are accurately labeled by domain experts and are now publicly available, scAlign can take advantage of the accurate labeling of these atlases to annotate new datasets that lack labels. Second, marker genes may be available for only a subset of cell types such as specific hematopoietic cells, in which case only a subset of cells may be reliably labeled. Even when marker genes are available, markers may not be unique to individual cell types and technical factors such as dropout may prevent truly expressed markers from being detected in the RNA. Here, scAlign can be used in conjunction with only the most confident labeled cells, or can even be used when there is overlapping labels (due to marker uncertainty).

Another advantage of scAlign over other integration methods is the improved ability to detect rare differential expression events between conditions. For typical alignment methods, once the effect of condition is removed via alignment, cells must still be clustered into putative cell types in order to identify which cells match across condition, and then perform an unpaired differential expression test within each cluster to identify condition-specific differences. The need to cluster cells means the detection of rare cell types can be highly sensitive to the choice of clustering algorithm or parameters. In contrast, through interpolation, scAlign predicts how each individual cell within the alignment space differs in expression between any of the input conditions, effectively performing a paired (or matched) differential expression calculation per-cell without the need to cluster. The result is scAlign can detect the presence of rare cell populations that differ in expression across conditions (**Fig. 9**).

scAlign implements two approaches to aligning more than two conditions simultaneously. In the reference-based alignment, a single reference condition is established and all other conditions are be aligned against the reference (**Additional file 1: Fig. S21**). This is expected to work well when all cell types are represented in all conditions, and has the benefit of speed. Alternatively, the all-pairs alignment mode performs an all-pairwise set of alignments simultaneously, which will be more robust to the presence of cell types only represented in a subset of the conditions.

The general design of scAlign’s neural network architecture and loss function makes it agnostic to the input RNA-seq data representation. Thus, the input data can either be gene-level counts, transformations of those gene level counts or the result of a preliminary step of dimensionality reduction such as principal component scores or canonical correlation vectors. In our study, we first transformed data into a relatively large number of principal component scores before input into scAlign, as this yielded much faster run times with little to no performance degradation. The improvement in computation time due to PCA pre-processing of the input data allowed scAlign to both converge more quickly and become feasible on a CPU-based system, therefore making scAlign a broadly applicable deep learning method. More generally, the design of scAlign’s neural network architecture and loss functions are general and not specific to scRNA-seq data. We therefore expect that scAlign should be applicable to any problem in which the study design consists of comparing two or more groups of unmatched samples, and where we expect there to be subpopulations of individuals within each group.

Here we have primarily compared scAlign against unsupervised alignment methods. In our supervised alignment results, scAlign compared favorably against the supervised methods MINT^1^ and scmap^37^ when assuming all cells are labeled. In the context of alignment, however, we reasoned that if a complete set of labels are available for all cells and conditions, then addressing the task of alignment is less useful, because cells of the same type across conditions can be directly compared via per-cell type differential expression analysis without alignment. Alternatively, in those contexts, each matching pair of cell types across conditions can be independently aligned using the unsupervised scAlign (or other unsupervised methods)_to identify matching subpopulations of cells.

The tasks of transcriptional alignment and batch correction of scRNA-seq data are intimately related, as one can view the biological condition of a cell as a batch whose effect should be removed before integrated data analysis. Compared to batch correction methods, scAlign leverages the flexibility of neural networks to perform alignment where cell states might exhibit heterogeneous responses to stimuli, yet through interpolation provides the interpretability that canonical batch correction methods enjoy.

Like all other supervised and unsupervised alignment methods, scAlign makes an underlying assumption that the two or more conditions used as input make sense biologically to align. That is, alignment methods assume that there are at least some common cell types between conditions that share some functional origin or similarity, that should be matched across conditions, even if they differ in state (e.g. expression) due to condition or stimulus. To the best of our knowledge, there is no procedure or strategy for identifying datasets that should not be aligned due to lack of matching cell types. As a result, any alignment method when applied to datasets which contain unrelated or dissimilar cell types can potentially lead to false positive matchings. This limitation is not specific to alignment methods; scRNA-seq analysis tools designed for other purposes, such as trajectory inference, assume that a trajectory exists in the input data in the first place, and will return a trajectory regardless of whether it makes sense to do so. scRNA-seq tools in general are useful for generating hypotheses (in the case of alignment, hypotheses about which cell types match across conditions, and how they differ), but need to be used cautiously by downstream users.

A related concern is the performance of alignment methods when there exists condition-specific cell types that have no matching cell type in another condition. In our experiments, we show that scAlign outperforms other alignment methods in this scenario by choosing a small number of cells from a matching cell type, and placing those small numbers of cells in the same region of alignment space as the condition-specific cell type; in other words, scAlign purposefully misaligns a small number of cells. scAlign tends to sacrifice a small number of cells because its objective function minimizes the distortion of the cell-cell pairwise similarity matrix within each input condition, and so sacrificing many cells would lead to a large distortion of the pairwise similarity matrix.

As a neural network-based method, scAlign usage requires specification of the network architecture before training, defined by the number of layers and number of nodes per layer. In our results, we have shown scAlign is largely robust to the size of the architecture, in part because in addition to the ridge penalty we apply to the weights of the network, our objective function minimizes the difference between the similarity matrix in the original expression and alignment spaces, which also acts as a form of data driven regularization.

## Declarations

### Ethics approval and consent to participate

Not applicable.

### Consent for publication

Not applicable.

### Availability of data and material

The code implementing scAlign, as well as the datasets generated and/or analysed in this study are available in the GitHub repository, at https://github.com/quon-titative-biology^48^ for all systems and on Bioconductor at http://bioconductor.org/packages/bioc/html/scAlign.html for Linux and Mac (starting November 2019). scAlign is released under Apache License 2.0.

### Authors’ contributions

NJJ and GQ conceived of the study, analyzed and interpreted the data, and wrote the manuscript. NJJ wrote the code for scAlign. All authors read and approved the final manuscript.

## Acknowledgements

We thank Bjorn Kafsack for assisting with interpretation of the P. falciparum data, and Manuel Llinás for pointing us to additional P. falciparum resources.

## Additional files

Additional file 1: Contains supplementary figures. (DOCX 11,959 KB)

Additional file 2: Contains supplementary gene ontology information. (XLS 50 KB)

Additional file 3: Contains supplementary marker gene information. (XLS 117 KB)

## Methods

### Methods overview

The scAlign method consists of two steps: (1) alignment, which learns a mapping from gene expression space of individual conditions into a common alignment space, and (2) interpolation, which learns a mapping from the common alignment space back to the gene expression space of the original conditions.

### Pairwise scRNA-seq alignment with scAlign

We define the alignment task as identifying a low dimensional embedding space (termed the alignment space) in which functionally similar cells map to the same coordinates. Viewed from the lens of perturbation studies, if sequencing a cell immediately before and after stimulus were possible, alignment would bring cells post-stimulus into the same region of alignment space as the cell before stimulus, therefore removing the effect of the stimulus.

scAlign encodes the alignment space by extending the recent approach of learning by association for neural networks^49,50^ into a unified framework for both unsupervised and supervised applications. For notational simplicity, we will assume we are aligning scRNA-seq data from a pair of conditions, though the framework extends to multiple conditions (see below). Let 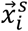 and 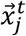 be vectors of length *G* that represent the gene expression profiles of cells *i* and *j* in conditions *s* and *t*, respectively. Similarly, let 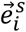 and 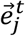 be vectors of length *K* that represent that alignment space embedding of cells *i* and *j* in conditions *s* and *t*, respectively, where the embeddings represent the linear activations of the final output layer of an encoder neural network.

scAlign trains an encoder neural network (parameterized by weights ***W***) that defines the alignment space by optimizing the network weights used to calculate 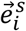 and 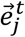 to minimize the following objective function:

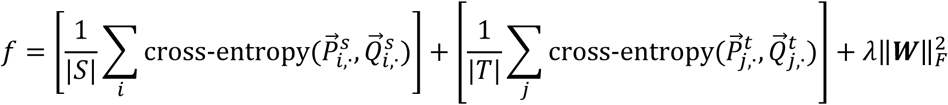

Where

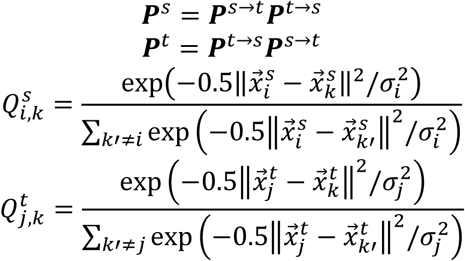

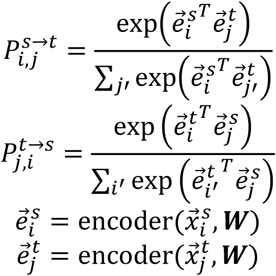

The central idea of the alignment procedure of scAlign is that it optimizes the embeddings of cells (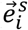 and 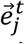) such that the scaled, pairwise cell-cell similarity matrix (or formally, a transition matrix) computed between cells within each condition in gene expression space (***Q***^***s***^ and ***Q***^***t***^) should be maintained within the alignment space (***P***^*s*^ and ***P***^*t*^), respectively. The novel aspect of scAlign compared to other dimensionality reduction methods is in how ***P***^*s*^ and ***P***^*t*^ are calculated. While ***P***^*s*^ would canonically be calculated by transforming the dot product of the embeddings 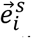 as is done in the tSNE method^51^ for example, scAlign computes roundtrip random walks of length two that traverse the two conditions. 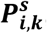, the transition probability of moving from cell *i* to cell *k* within condition *s*, is calculated as the probability of randomly walking from cell *i* to cell *k* in two steps: first from cell *i* to any cell *j* in the other condition *t* in the first step, then from that cell *j* to cell *k* (in condition *s*) in the second step. By forcing the random walk to first visit a cell in the other condition, scAlign encourages the encoder to bring cells from across the two conditions into similar regions of alignment space.

The network weights ***W*** are initialized by Xavier^52^ and optimized via the Adam algorithm^53^ with an initial learning rate of 10^−4^ and a maximum of 15,000 iterations. The neural network activation functions of each hidden layer are ReLU and the embedding layer has a linear activation function. Regularization is enforced through an L2 penalty on the weights along with per-layer batch normalization and dropout at a rate of 30%. The scAlign framework has three tunable parameters: the per-cell variance parameter 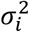 that controls the effective size of each cell’s neighborhood when defining the similarity matrix in gene expression space, the magnitude of the penalization term *λ* over ***W*** that is fixed at 10^−4^, and the size of the encoder network architecture.

For the tuning parameter 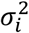, small values yield more local alignment, whereas larger values yield more global alignment. In our experiments, we train each model with a range of values for 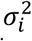. Typically, [5,10,30] provide robust results when training on mini-batches of less than 300 samples. While the per-cell variance parameter 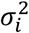 operates on the training mini-batch, we found training is robust to the choice of 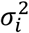.

In our experiments, we set the size of the encoder architecture by either automatically constructing a network based on the dimensionality of the input data in conjunction with a complexity parameter, or from a catalog of network architectures which are at most three layers deep. As with other neural networks, the size of the architecture defines the complexity and power of the network. Model complexity is important for alignment because the network must be powerful enough to align cells from conditions that yield heterogeneous responses to stimulus, but not so powerful that any cell in one condition can be mapped to any other cell in another condition, regardless of whether they are functionally related. We have found in our experiments (**Additional file 1: Fig. S3**) that the combination of cross-entropy loss and shrinkage applied to the network weights yields robustness to generously-large network architectures. Namely, by encouraging small weights and minimizing the differences in cell-cell similarity matrices between the expression and embedding spaces, we avoid training the neural network to perform unnecessary complex transformations on the data.

### Overview of multi-way alignment with scAlign

Alignment of three or more conditions simultaneously is implemented in two ways in the scAlign framework. In approach one (“all-pairs alignment”), round trip walks are computed between all pairs of conditions, and is expected to be the most accurate form of multi-way alignment. In approach two (‘reference-based alignment’), one condition is defined as a reference, against which all other conditions are aligned.

### All-pairs alignment with scAlign

In this strategy, we extend the pairwise alignment approach by performing round trip walks between all pairs of conditions simultaneously, while still sharing a single encoder’s neural network parameters across all conditions. Compared to the reference-based alignment approach below, the all-pairs approach will be more robust when there are cell types that are only represented in a subset of the input conditions. The objective function of the pairwise alignment approach is modified to include round trip walks between each condition *k* and the remaining conditions *l* ≠ *k*:

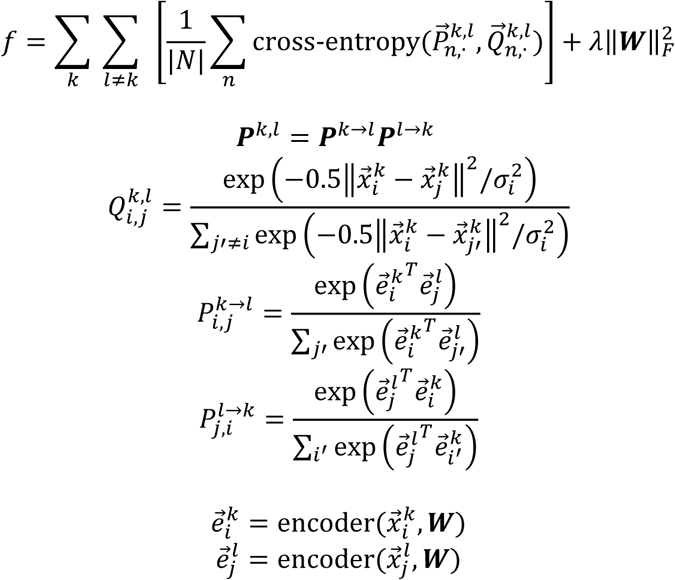

### Reference-based multi-way alignment with scAlign

In this strategy, multiple conditions are aligned simultaneously by selecting one condition to be a reference (*k*_ref_), against which all other conditions (*l* ≠ *k*_ref_) are aligned. Compared to the all-pairs approach, reference-based alignment is faster and therefore more scalable, though is expected to perform worse when there are cell types shared amongst non-reference conditions, that are not represented in the reference condition. The objective function for reference-based alignment is as follows:

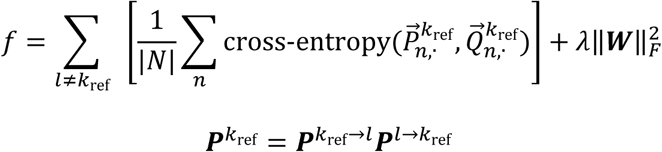

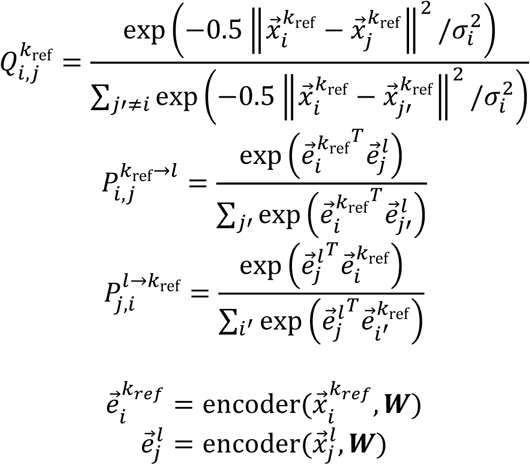

The remaining details for optimizing scAlign’s objective function in the multi-way case are identical to the paired alignment task described previously. We note that in our experiments the number of embedding dimensions had to be increased for three or more conditions in order to accommodate the increased information in the embeddings of the encoder shared across all *k* condtions.

### scRNA-seq interpolation with scAlign

The interpolation component of scAlign trains a condition-specific decoder to map cells from the alignment space back into each of the individual condition-specific gene expression spaces. The decoder network architecture is chosen to be symmetric with the encoder network trained during the alignment process, with weights randomly initialized and optimized again via the Adam optimizer^53^ with learning rate set at 10^−4^ and trained for at most 30,000 iterations.

### Calculation of the state variance map

After interpolating every cell (sequenced in any condition) from the alignment space back to every input condition, for each cell, we obtain multiple condition-specific representations for each cell. Then, per cell, we compute the variance of the interpolated expression patterns for that cell across the input conditions. The result is a matrix, termed the state variance map, which illustrates the variance in each gene-specific expression level for each cell predicted across conditions. In the special case where two conditions are being aligned, this state variance map can be viewed as a (predicted) paired differential expression map, where differences are calculated per cell.

### Shared autoencoder optimization

The training procedure for training a shared autoencoder followed that of scAlign in that the autoencoder was trained on data from all conditions simultaneously. The shared alignment space of the autoencoder was learned by optimizing with respect to the traditional mean squared error of reconstructing the original expression profiles for each condition by simultaneously training condition specific decoder networks.

### Principal Component Analysis and Canonical Correlation Analysis preprocessing transformations of scRNA-seq data

The objective function that scAlign optimizes does not incorporate terms specific to scRNA-seq data such as a negative binomial observation model. We found that computing the principal component and canonical correlates of the normalized scRNA-seq data and using the resulting scores in place of gene expression measurements maintained alignment and interpolation accuracy but sped up training significantly (**Additional file 1: Fig. S4**). Note that even when the encoder network is given PC or CC dimensions as input instead of gene expression measurements, the decoder is still trained to transform alignment space coordinates into the original gene expression space.

### Using partial or complete cell type labels with scAlign

The objective function optimized by scAlign can naturally incorporate partial, overlapping or complete cell type labels for the cells, in one or more conditions. Suppose there are *C* cell type labels available, in a pairwise alignment scenario. Then define matrix ***A***^*s*^ such that 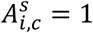 if cell *i* in condition *s* has cell type label *c*, else 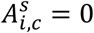. Similarly, define matrix *Â*^*s*^ containing the predicted class labels for all cells in condition *s*. The scAlign objective function then becomes:

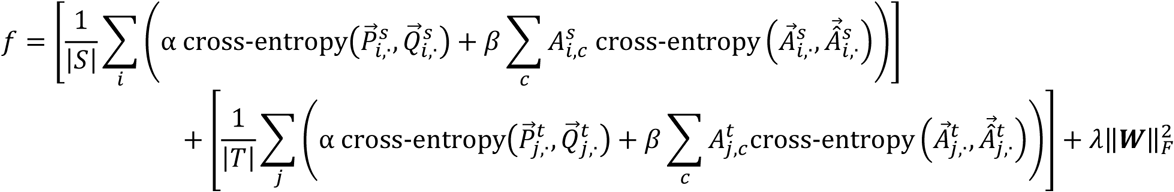

We incorporate partial, overlapping, or complete label information by introducing an extra set of terms corresponding to classification loss and weighted by the factor *β*. The classifier loss terms minimize the mean cross-entropy of the predicted and actual cell labels as defined by the second term within each summation of *f*. The adaptation and classifier components *f* are balanced by hyperparameter weights *α* and *β* respectively. Adjusting *α* and *β* allows emphasis to be placed individually on the pairwise cell similarity or known labels; in this work both weights were fixed to 1.0 when label information is provided.

### Acquisition and preprocessing of Mann et al. benchmark

We obtained the gene count matrix for HSC data generated from Mann et al.^40^ from GSE100426. The provided data matrix was already filtered based on quality control metrics. We normalized the count matrix to TP10K and then removed plate specific batch effects by fitting a linear model on the scaled and centered data using Seurat’s NormalizeData and ScaleData functions. We retained the union of the top 3,000 variable genes between control and condition cells.

### Acquisition and preprocessing of Kowalczyk et al. benchmark

We obtained the gene count matrix for both C57BL6 and DBA mouse HSC data generated from Kowalczyk et al.^39^ from GSE59114. Only single cell data from mouse C57BL6 was used during alignment to avoid cross mouse batch effects. We normalized the count matrix to TP10K then scaled and centered using Seurat’s NormalizeData and ScaleData functions. We retained the union of the top 3,000 variable genes between young and old cells and genes associated with GO terms reported in Kowalczyk et al.

### Acquisition and preprocessing of CellBench benchmark

We obtained the gene count matrix for the RNA mixture experiments in CellBench generated by Tian et al.^38^ from the R data file mRNAmix_qc.RData available on GitHub. We normalized the count matrix to TP10K then scaled and centered using Seurat’s NormalizeData and ScaleData functions. We retained the union of the top 3,000 variable genes between mixtures profiled on CEL-Seq2 and SORT-Seq.

### Execution of scAlign for benchmark data

We provided scAlign with normalized and scaled gene expression following standard Seurat preprocessing protocols. The most variable genes were identified using FindVariableGenes function implemented in Seurat which was used to subset the data matrices. scAlign was then trained with default parameter settings including 15,000 steps, mini batch size of 150, perplexity of 30, a 3-layer neural network with 32 dimensions in the final embedding layer.

### Execution of scAlign for *P. falciparum* (malaria parasite) data

We provided scAlign with the top 26 PCs as reported by Poran et al. and available on the Kafsack lab Github. scAlign was then trained for 15,000 steps, mini batch size of 1000, perplexity of 100, a 3-layer neural network with 32 dimensions in the final embedding layer.

### Execution of scAlign for pancreatic islet data

We provided scAlign with the top 30 canocical correlation vectors as computed by Seurat following the standard preprocessing pipeline. scAlign was then trained with default parameter settings primarily defined by 15,000 steps, mini batch size of 1000, perplexity of 100, a 3-layer neural network with 64 dimensions in the final embedding layer.

### Execution of other scRNA-seq alignment methods

We compared scAlign against MNN^2^, Seurat^3^, scMerge^54^, Scanorama^2^, scVI^7^, MINT^1^ and scmap^5^. Each method was run based on method-specific guidelines provided by the original authors and following the workflow defined by CellBench publicly available on GitHub. Prior to running each method, the FindVariableGenes function implemented in Seurat was used to identify the most variable genes for a consistent subsetting of the following data matrices. MNN was provided log-count data subset to the most variable genes with all parameters set to default. Seurat v2 and v3 were provided the count level data which was normalized, then scaled and centered using the NormalizeData and ScaleData functions. Initially, 30 canonical correlates were used for dimensionality reduction, then the MetageneBicorPlot function was used to select the optimal number of dimensions as defined by Seurat’s integrated PBMC tutorial. The remaining canonical correlates were aligned using the Seurat v2 AlignSubspace function or Seurat v3 FindIntegrationAnchors and IntegrateData functions. scMerge was provided both count and log-count data along with a set of least variable genes identified by sorting the results of the var function in R on the normalized count matrix. The parameter kmeansK that specifies the number of clusters was set based on cell type information. Supervised scMerge (scMerge+) was additionally provided cell labels and the “cell_type_match” parameter was set to true. Scanorama was provided with log-count data subset to the most variable genes previously identified by decompseVar, and return_dense was set to TRUE. scVI was provided with the full count data and trained until convergence based on log likelihood for the best model parameterization identified by grid search (see below). MINT was provided with the log-count data and the cell type labels. scmap was provided the count and log-count data and both indexCluster and indexCell functions were used to compute cross condition labels.

### Execution of scVI and parameter search

The provided tutorials for scVI varied the input parameters to scVI and did not provide further guidance how to select parameters. We therefore performed a grid search over scVI parameters, and chose the parameter combination that minimized the reported loss. The grid search was performed with respect to number of layers (1, 2 or 3), number of epochs (1000, 2500 or 5000) and learning rate (0.01, 0.001, or 0.0001) **(Additional file 1: Fig. S22).**

### Construction of HeterogeneousBenchmark

We constructed the HeterogeneousBenchmark benchmark by merging multiple count matrices from the Mann et al. and Kowalczyk et al. studies. The control condition was defined completely by young C57BL/6 mouse cells. To construct the stimulated condition, we merged LT-HSCs perturbed by LPS from Mann et al., ST-HSCs from old C57BL/6 mouse cells and MPPs from both young and old DBA mouse cells collected by Kowalczyk et al.

### Acquisition and preprocessing of *P. falciparum* (malaria parasite) data

We obtained the gene count matrix for the *P. falciparum* data generated by Poran et al.^45^ from the KafsackLab GitHub repository. The data was preprocessed using the provided scripts and subset into the +/−Shld conditions using the metadata.

### Acquisition and preprocessing of pancreatic islet data

We obtained the gene count matrix for the human pancreatic islet datasets from the following accessions: GSE81076 (CEL-Seq), GSE85241 (CEL-Seq2), GSE86469 (Fluidigm C1), and E-MTAB-5061 (Smart-Seq2). Following preprocessing previously defined by Stuart et al.^35^, we filtered out cells for which fewer than 1,750 unique genes/cell (CEL-Seq) or 2,500 genes/cell (CEL-Seq2/Fluidigm C1/Smart-Seq2) were detected.

### Identification of differentially expressed genes

Differentially expressed (DE) genes were computed using the bimod, DESeq2 and MAST methods implemented in the Seurat findMarkers function. The intersection of DE genes with P-value less than 0.01 from these three methods was used to define a final set of DE genes for each cell type. The analysis was performed on the normalized, scaled and centered data matrices computed by Seurat’s preprocessing pipeline.

### Identification of robust marker genes

Differentially expressed (DE) genes were computed using bimod^55^, DESeq2^56^ and MAST^57^ methods implemented in Seurat findMarkers function for each cell type individually for each condition. The union of cell type marker genes with corrected p-value less than 0.05 was used to define a final set of marker genes for each condition. Additionally, genes which were highly correlated (>0.9) with the condition-specific marker genes were also included. Finally, we took the intersection of these marker gene sets to define the robust common marker genes across conditions. The analysis was performed on the normalized, scaled and centered data matrices computed by Seurat’s preprocessing pipeline.

### Construction of matched gene sets

We first computed the mean expression level of all genes, and created five approximately equally sized bins representing groups of genes with lowest to highest expression. For the robust common marker gene set, we identified the number of common marker genes that came from each bin. We then created control gene sets by drawing random sets of genes of the same size as the robust common marker set, and with the same distribution of genes over the five bins. We repeated the sampling procedure 10,000 times to obtain a representative collection of matched gene sets.

### Measuring the deviation in cell embeddings by *in silico* gene set perturbation

To determine the importance of a single gene set to scAlign’s calculation of the embedded representation of each cell, we zeroed out the expression measurements of all genes in the gene set across all cells. We then measure the median and maximum change (using Euclidean distance) in cell embeddings before and after zero-ing out the expression measurements. To compute a P-value, we generated random gene sets of the same size and matched for expression levels of the genes, and calculated the number of random gene sets that yielded a deviation at least as large as what we observed for a gene set of interest.

### Measuring accuracy of pairwise alignments

Alignment performance for each method was measured as a weighted combination of cross-condition label prediction accuracy and alignment score^3^. The cross-condition label prediction was performed by training a classifier to label one condition (stimulated condition by default) using only labels from the corresponding control condition. Specifically, a K-nearest neighbors classifier from the R library ‘class’ was initialized with control cell embeddings after alignment, along with their corresponding cell type labels. The classifier was then used to predict labels for the stimulated cells. The predicted labels were compared against heldout labels to measure accuracy. The final score accuracy_composite_ is defined by the product of the classifier accuracy and alignment score.

### Measuring accuracy of multi-way (three or more) alignments

Similarly, to measure alignment performance on the alignment of three or more conditions, we measured the weighted combination of a representative-based label prediction accuracy and alignment score. The representative-based label prediction was performed by iteratively treating each condition as the representative, and training a classifier to label cells from all non-representative conditions using only labels and cells from the single representative condition. The mean accuracy was computed for all condition specific label predictions as the final accuracy. As a classifier, we chose a K-nearest neighbors classifier from the R library ‘class’ and initialized it with the representative condition cell embeddings after alignment, along with their corresponding cell type labels. The classifier was then used to predict labels for all the non-representative cells. The predicted labels were compared against heldout labels to measure accuracy. The final score accuracy_composite_ is defined by the product of the mean accuracy and alignment score.

### Measuring accuracy of transcriptional interpolation

To measure interpolation accuracy, we measured the ability of a classifier trained on the gene expression data of the cells measured under one condition to correctly label interpolated gene expression profiles of cells sequenced under the other condition (but interpolated into the current condition). A K-nearest neighbors classifier from the R library ‘class’ was initialized with 90% of expression data and tested on the remaining heldout set of 10% to define gene expression specific accuracy. The classifier was then used to predict the labels for cells represented by interpolated gene expression values to compute an interpolation specific accuracy. 10-fold cross validation was performed using this procedure and the average accuracy was reported.

### 2D tSNE visualizations of embeddings for alignment methods

By default, we use the Rtsne implementation of tSNE, which first projects input data into 50 principal components before inputting into the tSNE algorithm. All methods other than Seurat and scAlign produce corrected expression matrices, and for these we use the default 50 PCs for Rtsne. Seurat automatically selects the number of dimensions to project into for each individual condition. scAlign was used to align scRNA-seq data into a 32-dimensional embedding space for all runs. For both Seurat and scAlign, the PCA step of Rtsne was skipped.

## List of Abbreviations

scRNA-seq: Single-cell RNA sequencing
LT-HSC: Long-term hematopoietic stem cell
ST-HSC: Short-term hematopoietic stem cell
MPP: Multi-potent progenitor
DEG: differentially expressed gene
LPS: Lipopolysaccharide
PCA: Principal components analysis
CCA: Canonical correlation analysis
tSNE: t-distributed stochastic neighbor embedding
UMAP: Uniform Manifold Approximation and Projection

## References

1. Rohart, F., Eslami, A., Matigian, N., Bougeard, S. & Lê Cao, K.-A. MINT: a multivariate integrative method to identify reproducible molecular signatures across independent experiments and platforms. BMC Bioinformatics 18, 128 (2017).

2. Haghverdi, L., Lun, A. T. L., Morgan, M. D. & Marioni, J. C. Batch effects in single-cell RNA-sequencing data are corrected by matching mutual nearest neighbors. Nat. Biotechnol. 36, 421–427 (2018).

3. Butler, A., Hoffman, P., Smibert, P., Papalexi, E. & Satija, R. Integrating single-cell transcriptomic data across different conditions, technologies, and species. Nat. Biotechnol. 36, 411–420 (2018).

4. Lin, Y. et al. scMerge leverages factor analysis, stable expression, and pseudoreplication to merge multiple single-cell RNA-seq datasets. Proc. Natl. Acad. Sci. 201820006 (2019). doi:10.1073/pnas.1820006116

5. Kiselev, V. Y., Yiu, A. & Hemberg, M. scmap: projection of single-cell RNA-seq data across data sets. Nat. Methods 15, 359–362 (2018).

6. Argelaguet, R. et al. Multi-Omics factor analysis - a framework for unsupervised integration of multi-omic data sets. bioRxiv 217554 (2018). doi:10.1101/217554

7. Lopez, R., Regier, J., Cole, M. B., Jordan, M. I. & Yosef, N. Deep generative modeling for single-cell transcriptomics. Nat. Methods 15, 1053–1058 (2018).

8. Hie, B. L., Bryson, B. & Berger, B. Panoramic stitching of heterogeneous single-cell transcriptomic data. bioRxiv (2018). doi:10.1101/371179

9. Kiselev, V. Y. et al. SC3: consensus clustering of single-cell RNA-seq data. Nat. Methods 14, 483–486 (2017).

10. Ji, Z. & Ji, H. TSCAN: Pseudo-time reconstruction and evaluation in single-cell RNA-seq analysis. Nucleic Acids Res. 44, e117–e117 (2016).

11. Trapnell, C. et al. The dynamics and regulators of cell fate decisions are revealed by pseudotemporal ordering of single cells. Nat. Biotechnol. 32, 381–386 (2014).

12. Qiu, X. et al. Reversed graph embedding resolves complex single-cell trajectories. Nat. Methods 14, 979–982 (2017).

13. Risso, D., Ngai, J., Speed, T. P. & Dudoit, S. Normalization of RNA-seq data using factor analysis of control genes or samples. Nat. Biotechnol. 32, 896–902 (2014).

14. Lawlor, N. et al. Single-cell transcriptomes identify human islet cell signatures and reveal cell-type-specific expression changes in type 2 diabetes. Genome Res. 27, 208–222 (2017).

15. Muraro, M. J. et al. A Single-Cell Transcriptome Atlas of the Human Pancreas. Cell Syst. 3, 385–394.e3 (2016).

16. Segerstolpe, Å. et al. Single-Cell Transcriptome Profiling of Human Pancreatic Islets in Health and Type 2 Diabetes. Cell Metab. 24, 593–607 (2016).

17. Büttner, M., Miao, Z., Wolf, F. A., Teichmann, S. A. & Theis, F. J. A test metric for assessing single-cell RNA-seq batch correction. Nat. Methods 16, 43 (2019).

18. Stuart, T. & Satija, R. Integrative single-cell analysis. Nat. Rev. Genet. 20, 257 (2019).

19. Subramanian, A. et al. A Next Generation Connectivity Map: L1000 Platform and the First 1,000,000 Profiles. Cell 171, 1437–1452.e17 (2017).

20. Datlinger, P. et al. Pooled CRISPR screening with single-cell transcriptome readout. Nat. Methods 14, 297–301 (2017).

21. Dixit, A. et al. Perturb-Seq: Dissecting Molecular Circuits with Scalable Single-Cell RNA Profiling of Pooled Genetic Screens. Cell 167, 1853–1866.e17 (2016).

22. Jaitin, D. A. et al. Dissecting Immune Circuits by Linking CRISPR-Pooled Screens with Single-Cell RNA-Seq. Cell 167, 1883–1896.e15 (2016).

23. Hodge, R. D. et al. Conserved cell types with divergent features between human and mouse cortex. bioRxiv 384826 (2018). doi:10.1101/384826

24. Hon, C.-C., Shin, J. W., Carninci, P. & Stubbington, M. J. T. The Human Cell Atlas: Technical approaches and challenges. Brief. Funct. Genomics 17, 283–294 (2018).

25. Tabula Muris Consortium et al. Single-cell transcriptomics of 20 mouse organs creates a Tabula Muris. Nature 562, 367–372 (2018).

26. Vento-Tormo, R. et al. Single-cell reconstruction of the early maternal-fetal interface in humans. Nature 563, 347–353 (2018).

27. Moffitt, J. R. et al. Molecular, spatial, and functional single-cell profiling of the hypothalamic preoptic region. Science 362, (2018).

28. Hodge, R. D. et al. Conserved cell types with divergent features between human and mouse cortex. bioRxiv 384826 (2018). doi:10.1101/384826

29. Plasschaert, L. W. et al. A single-cell atlas of the airway epithelium reveals the CFTR-rich pulmonary ionocyte. Nature 560, 377–381 (2018).

30. Hicks, S. C., Townes, F. W., Teng, M. & Irizarry, R. A. Missing data and technical variability in single-cell RNA-sequencing experiments. Biostat. Oxf. Engl. 19, 562–578 (2018).

31. Buettner, F. et al. Computational analysis of cell-to-cell heterogeneity in single-cell RNA-sequencing data reveals hidden subpopulations of cells. Nat. Biotechnol. 33, 155–160 (2015).

32. Shalek, A. K. et al. Single-cell RNA-seq reveals dynamic paracrine control of cellular variation. Nature 510, 363–369 (2014).

33. Eldar, A. & Elowitz, M. B. Functional roles for noise in genetic circuits. Nature 467, 167–173 (2010).

34. Maamar, H., Raj, A. & Dubnau, D. Noise in gene expression determines cell fate in Bacillus subtilis. Science 317, 526–529 (2007).

35. Stuart, T. et al. Comprehensive integration of single cell data. bioRxiv 460147 (2018). doi:10.1101/460147

36. Haghverdi, L., Lun, A. T. L., Morgan, M. D. & Marioni, J. C. Batch effects in single-cell RNA-sequencing data are corrected by matching mutual nearest neighbors. Nat. Biotechnol. 36, 421–427 (2018).

37. Kiselev, V. Y., Yiu, A. & Hemberg, M. scmap: projection of single-cell RNA-seq data across data sets. Nat. Methods 15, 359–362 (2018).

38. Tian, L. et al. scRNA-seq mixology: towards better benchmarking of single cell RNA-seq protocols and analysis methods. bioRxiv 433102 (2018). doi:10.1101/433102

39. Kowalczyk, M. S. et al. Single-cell RNA-seq reveals changes in cell cycle and differentiation programs upon aging of hematopoietic stem cells. Genome Res. 25, 1860–1872 (2015).

40. Mann, M. et al. Heterogeneous Responses of Hematopoietic Stem Cells to Inflammatory Stimuli are Altered with Age. bioRxiv 163402 (2017). doi:10.1101/163402

41. Trapnell, C. Defining cell types and states with single-cell genomics. Genome Res. 25, 1491–1498 (2015).

42. Ji, Z. & Ji, H. TSCAN: Pseudo-time reconstruction and evaluation in single-cell RNA-seq analysis. Nucleic Acids Res. gkw430 (2016). doi:10.1093/nar/gkw430

43. Setty, M. et al. Wishbone identifies bifurcating developmental trajectories from single-cell data. Nat. Biotechnol. 34, 637–645 (2016).

44. Welch, J. D., Hartemink, A. J. & Prins, J. F. SLICER: inferring branched, nonlinear cellular trajectories from single cell RNA-seq data. Genome Biol. 17, 106 (2016).

45. Poran, A. et al. Single-cell RNA sequencing reveals a signature of sexual commitment in malaria parasites. Nature advance online publication, (2017).

46. Bancells, C. et al. Revisiting the initial steps of sexual development in the malaria parasite Plasmodium falciparum. Nat. Microbiol. 4, 144–154 (2019).

47. Josling, G. A. et al. Regulation of sexual differentiation is linked to invasion in malaria parasites. (Microbiology, 2019). doi:10.1101/533877

48. Johansen, N.J., Quon, G. scAlign R code. Zenodo. https://doi.org/10.5281/zenodo.3339657.

49. Haeusser, P., Mordvintsev, A. & Cremers, D. Learning by Association - A versatile semi-supervised training method for neural networks. in IEEE Conference on Computer Vision and Pattern Recognition (CVPR) (2017).

50. Haeusser, P., Frerix, T., Mordvintsev, A. & Cremers, D. Associative Domain Adaptation. in IEEE International Conference on Computer Vision (ICCV) (2017).

51. Maaten, L. van der & Hinton, G. Visualizing Data using t-SNE. J. Mach. Learn. Res. 9, 2579–2605 (2008).

52. Glorot, X. & Bengio, Y. Understanding the difficulty of training deep feedforward neural networks. in Proceedings of the Thirteenth International Conference on Artificial Intelligence and Statistics (eds. Teh, Y. W. & Titterington, M.) 9, 249–256 (PMLR, 2010).

53. Kingma, D. P. & Ba, J. Adam: A Method for Stochastic Optimization. ArXiv14126980 Cs (2014).

54. Lin, Y. et al. scMerge: Integration of multiple single-cell transcriptomics datasets leveraging stable expression and pseudo-replication. bioRxiv 393280 (2018). doi:10.1101/393280

55. McDavid, A. et al. Data exploration, quality control and testing in single-cell qPCR-based gene expression experiments. Bioinforma. Oxf. Engl. 29, 461–467 (2013).

56. Love, M. I., Huber, W. & Anders, S. Moderated estimation of fold change and dispersion for RNA-seq data with DESeq2. Genome Biol. 15, 550 (2014).

57. Finak, G. et al. MAST: a flexible statistical framework for assessing transcriptional changes and characterizing heterogeneity in single-cell RNA sequencing data. Genome Biol. 16, 278 (2015).

