## Supplemental Figures for "scAlign: a tool for alignment, integration and rare cell identification from scRNA-seq data"

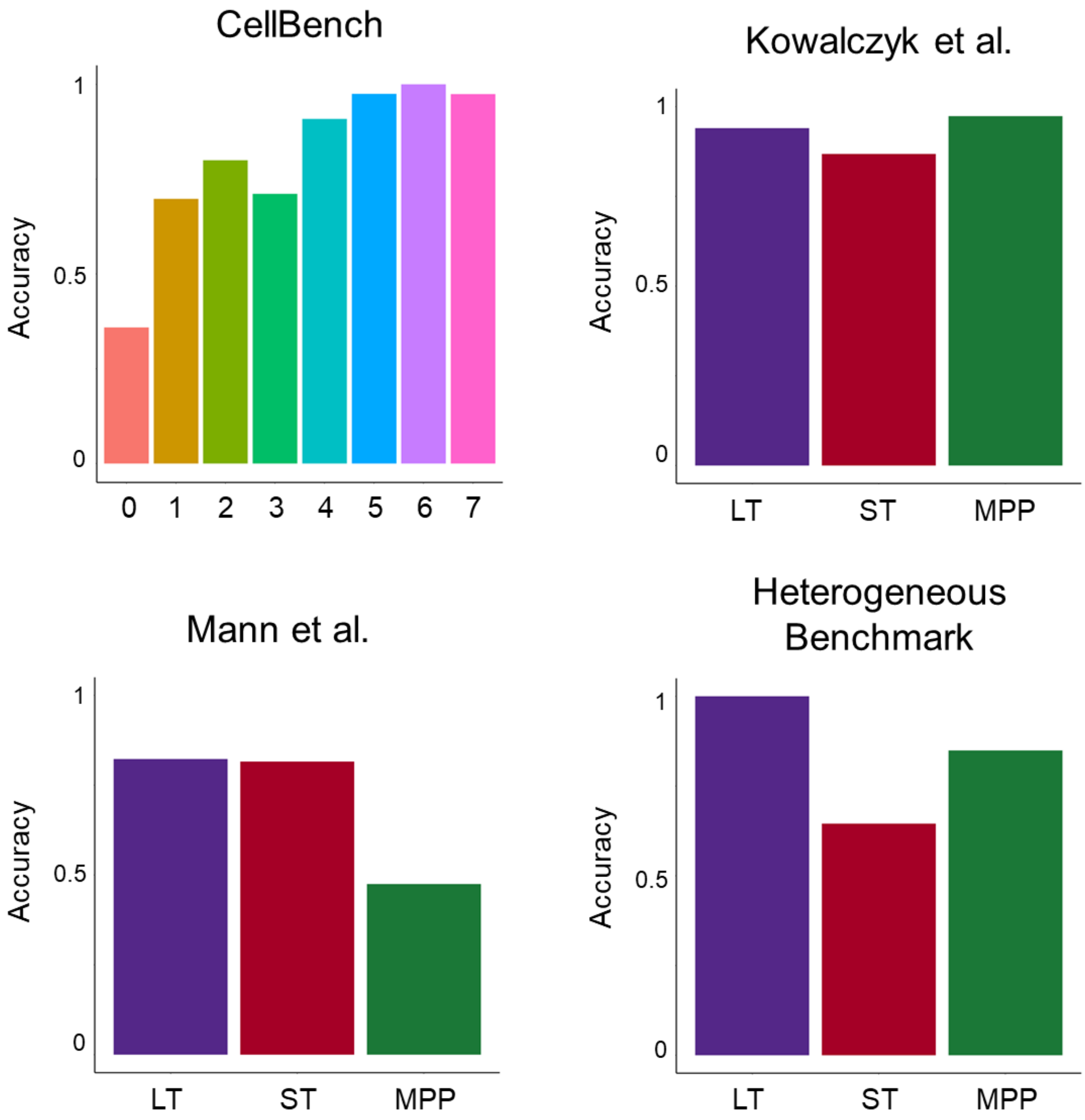

**Supplementary Figure 1: Cell type-specific accuracy for each of the four benchmark datasets.** Bar plots indicate the accuracy with respect to each cell type.

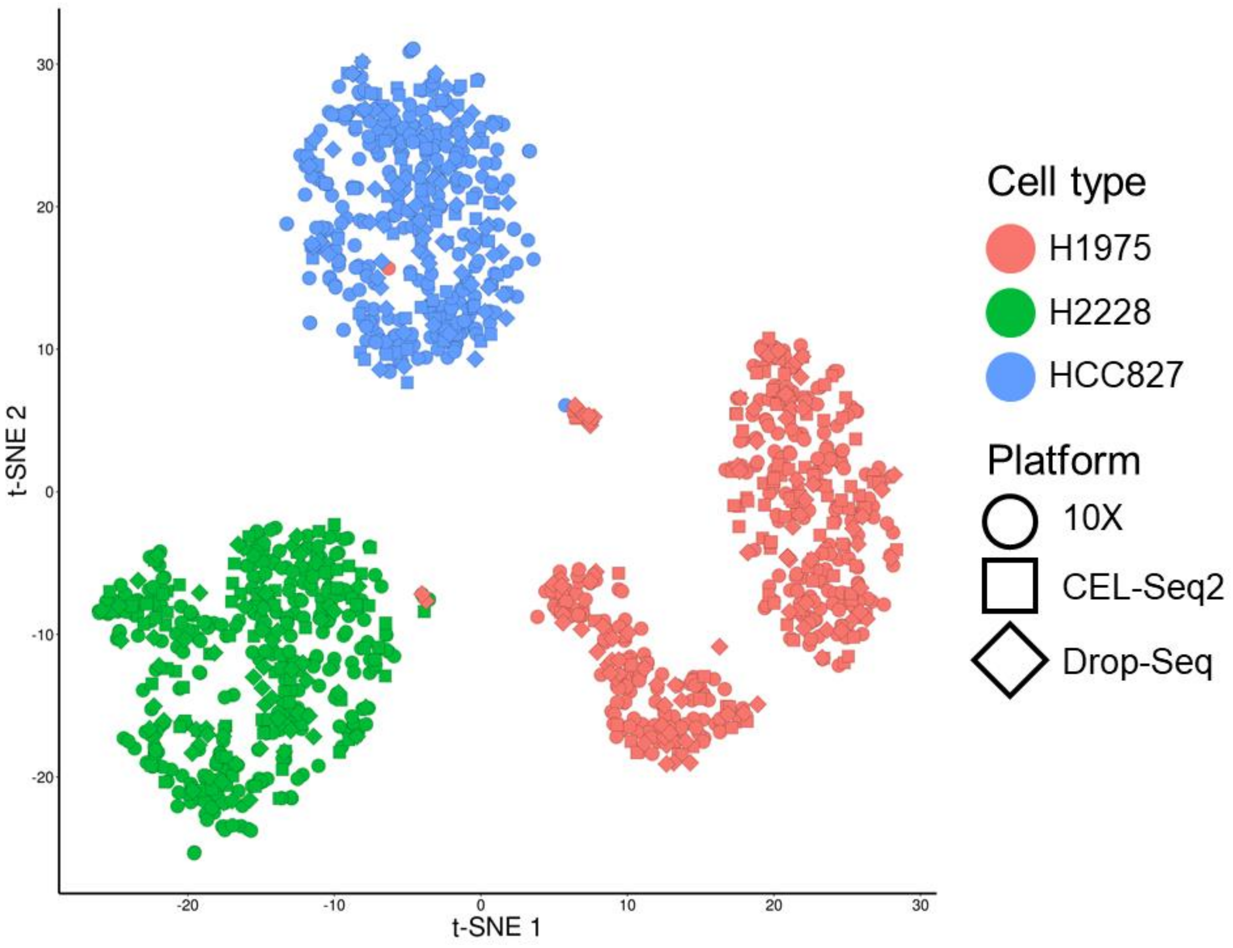

**Supplementary Figure 2: Standard normalization procedures align the same cell types sequenced using different protocols by CellBench.** Scatterplot of the three human lung adenocarcinoma cell lines sequenced by CellBench using either 10X Chromium, CEL-Seq2 or Drop-Seq, and normalized independently using Seurat.

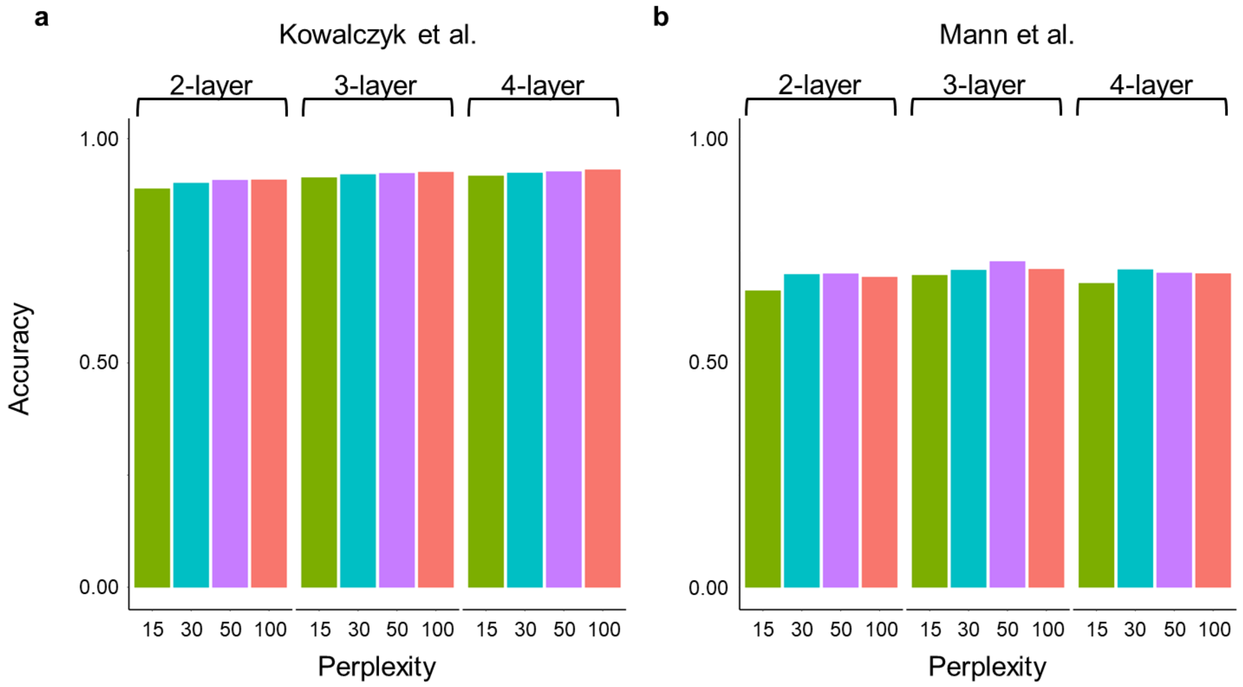

**Supplementary Figure 3: Robustness of scAlign to neural network architecture and size**. Barplots indicate accuracy of different network architectures, hidden layers, and perplexity parameter settings (x-axis) for scAlign trained on the Kowalcyk et al. and Mann et al. benchmarks.

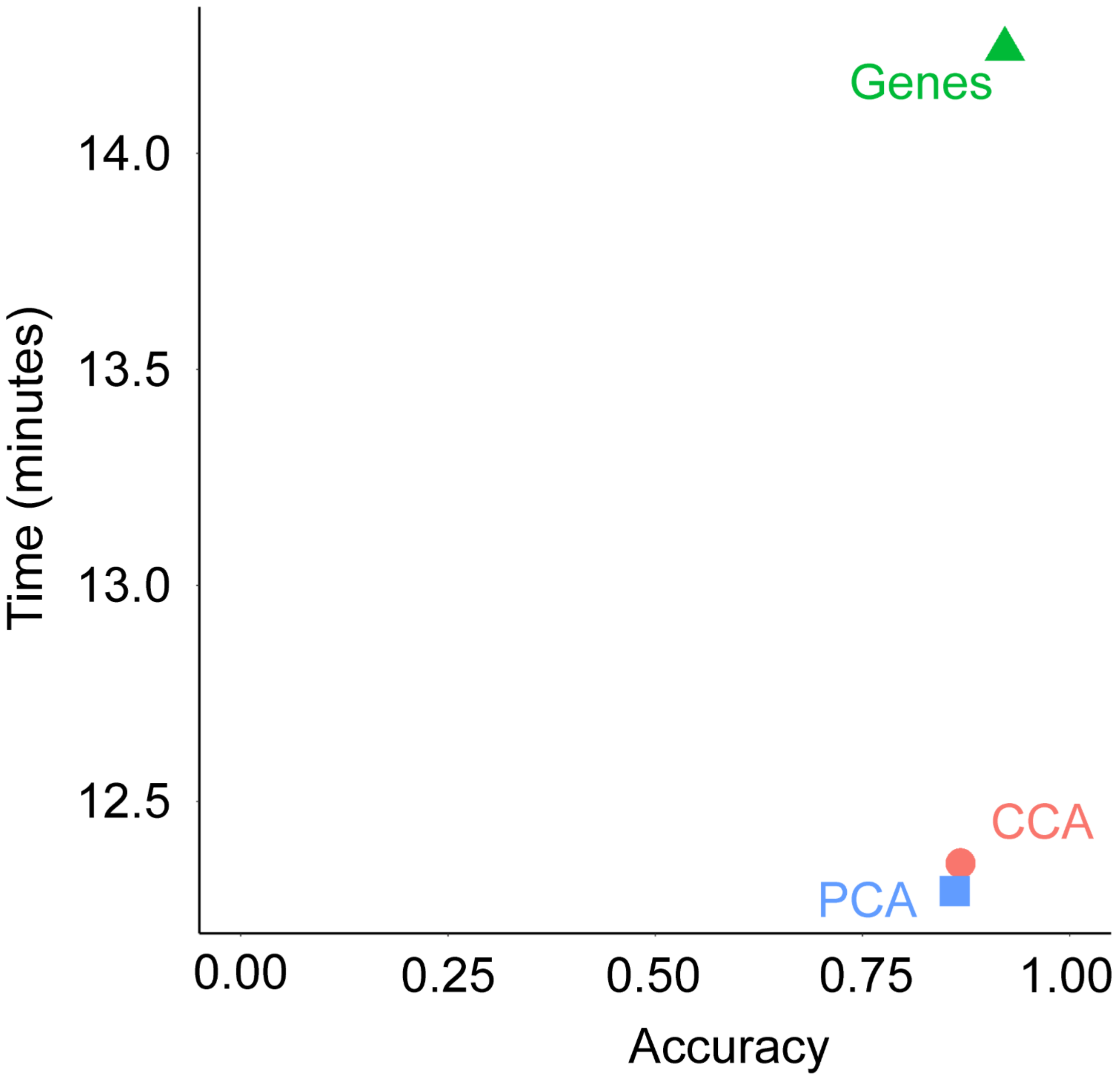

**Supplementary Figure 4: Initial data preprocessing step of dimensionality reduction increases computation speed without degrading accuracy or alignment.** Scatterplot indicates the computation time on the y-axis and cross condition classification accuracy on the x-axis after training scAlign with the top 3,000 variant genes, 10 CCs or 20 PCs.

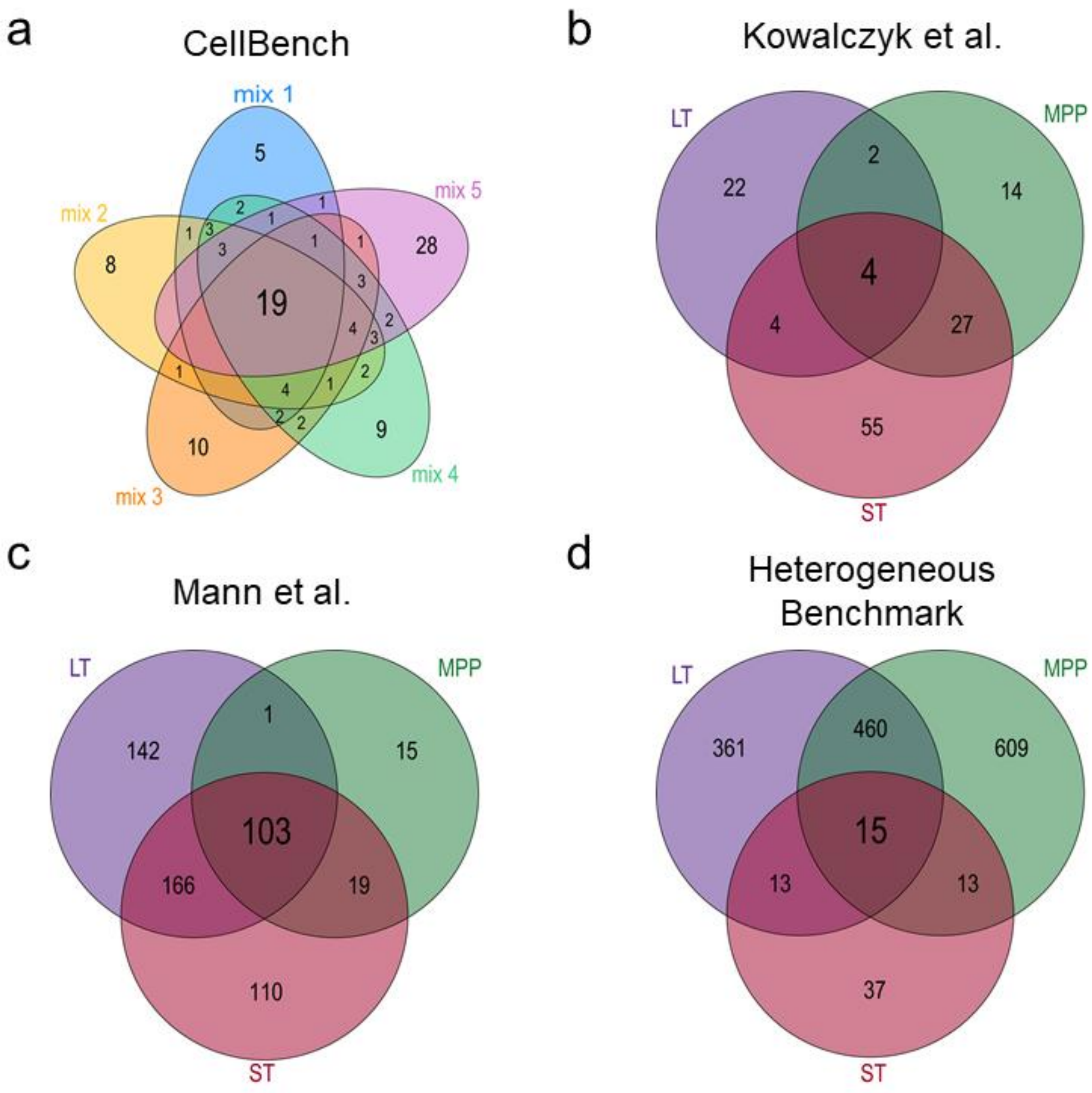

**Supplementary Figure 5: Overlap of differentially expressed genes for the four benchmark datasets. (a-d)** Venn diagram indicating the number of overlapping DEGs (measured across condition) for each cell type in the four benchmarks.

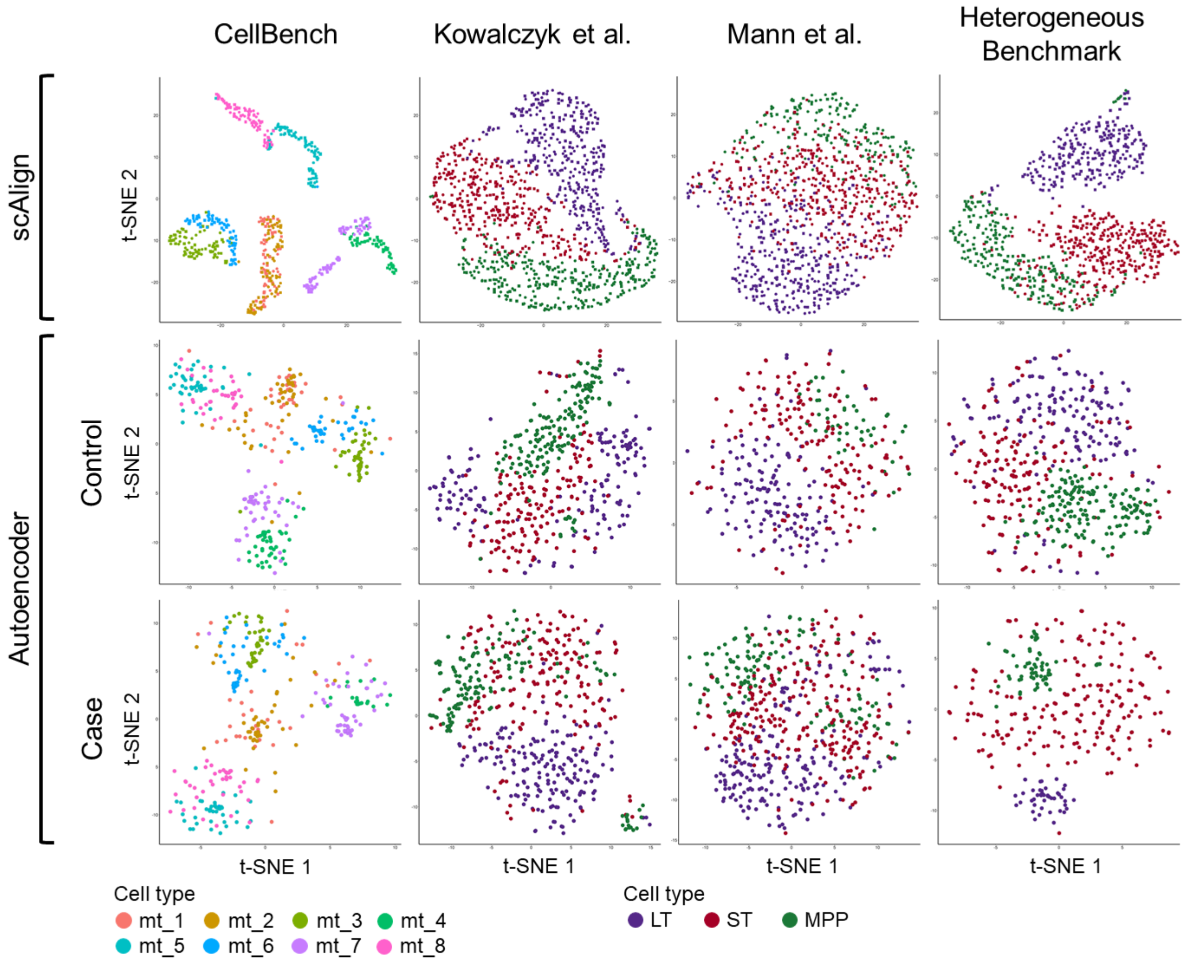

**Supplementary Figure 6: Comparison of the embeddings learned by scAlign and an autoencoder.** t-SNE visualization of the embeddings after training either scAlign or a condition-specific autoencoder on each of the four benchmarks.

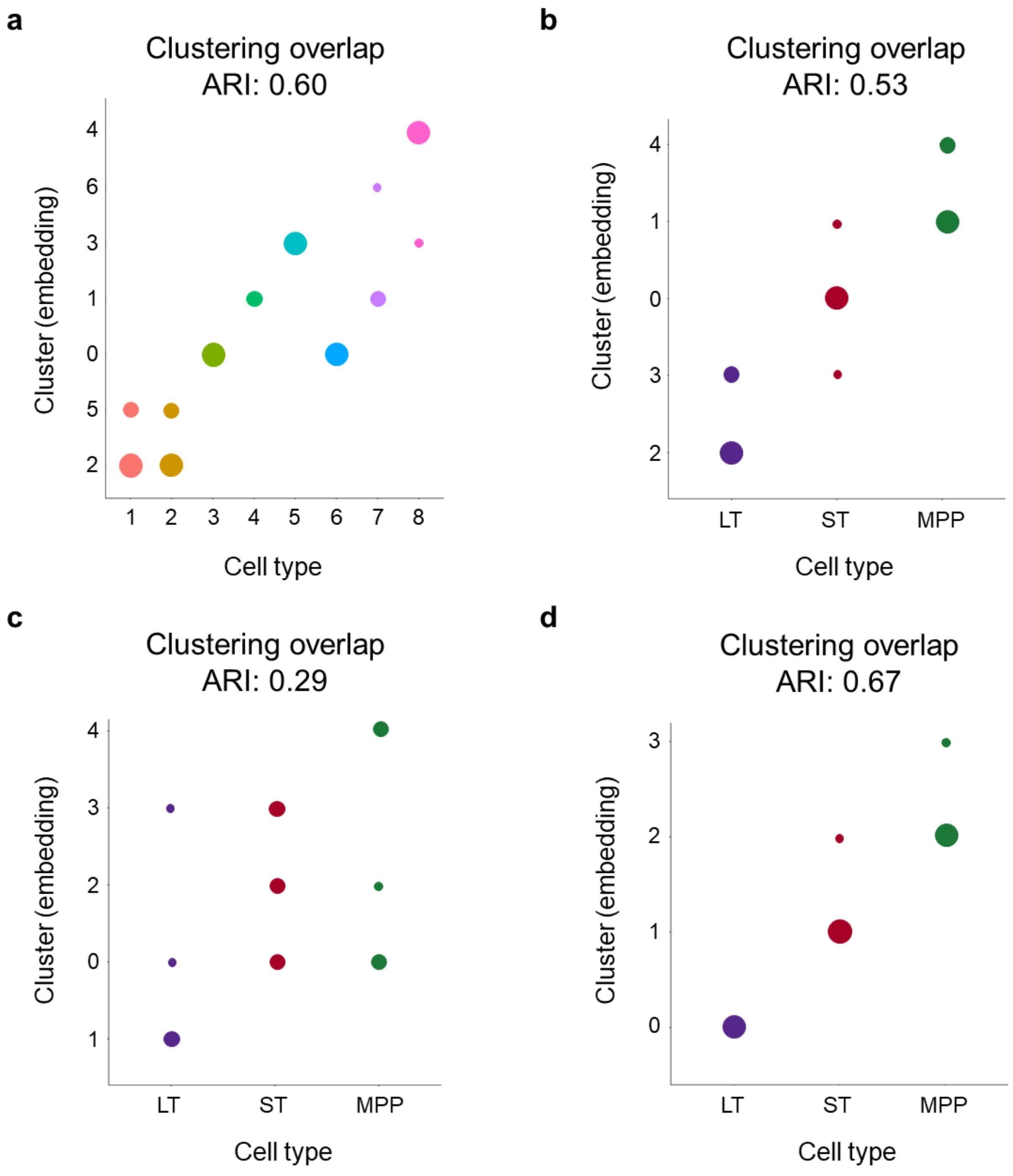

**Supplementary Figure 7: Comparison of clustering in scAlign's alignment space and known cell type labels.** (**a**) Dot plot visualizes the amount of overlap between de novo clustering and cell type annotations for CellBench. (b-d) Similarly to (a) but for Kowalczyk et al., Mann et al. and HeterogeneousBenchmark, respectively.

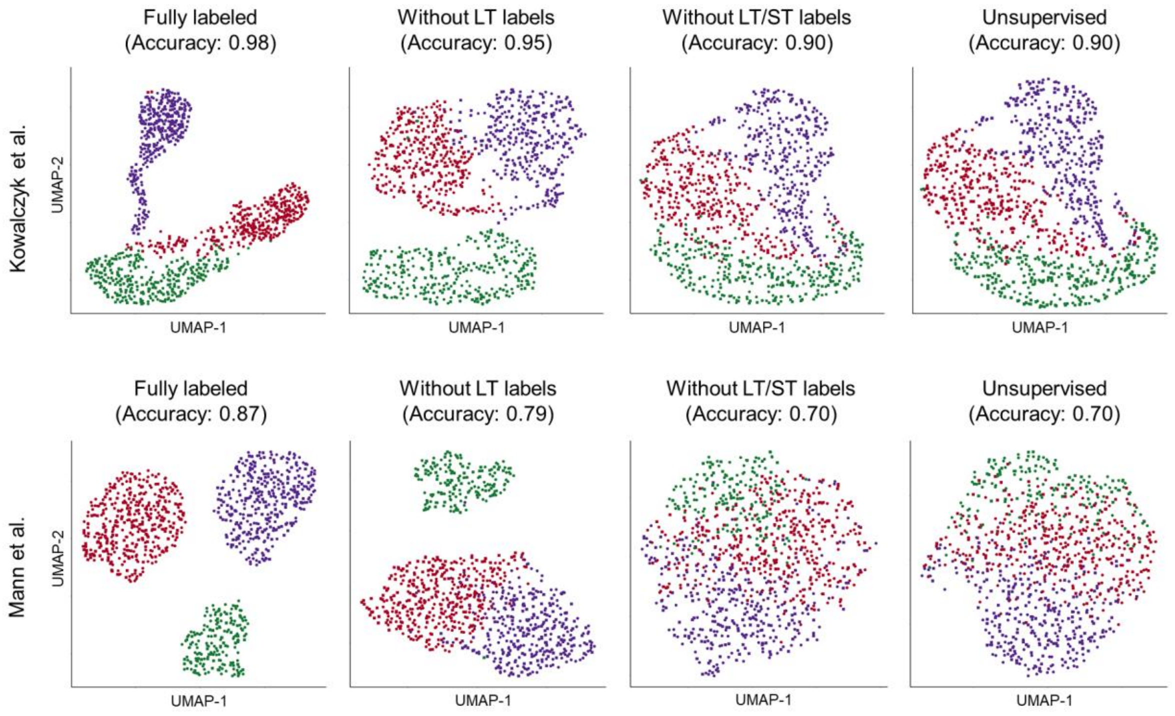

**Supplementary Figure 8: Partial labels yield performance between fully labeled and no-label data.** UMAP visualization of scAlign alignment of Kowalczyk et al. and Mann et al. after training with complete cell label information or after removing either the LT or LT and ST condition specific cells.

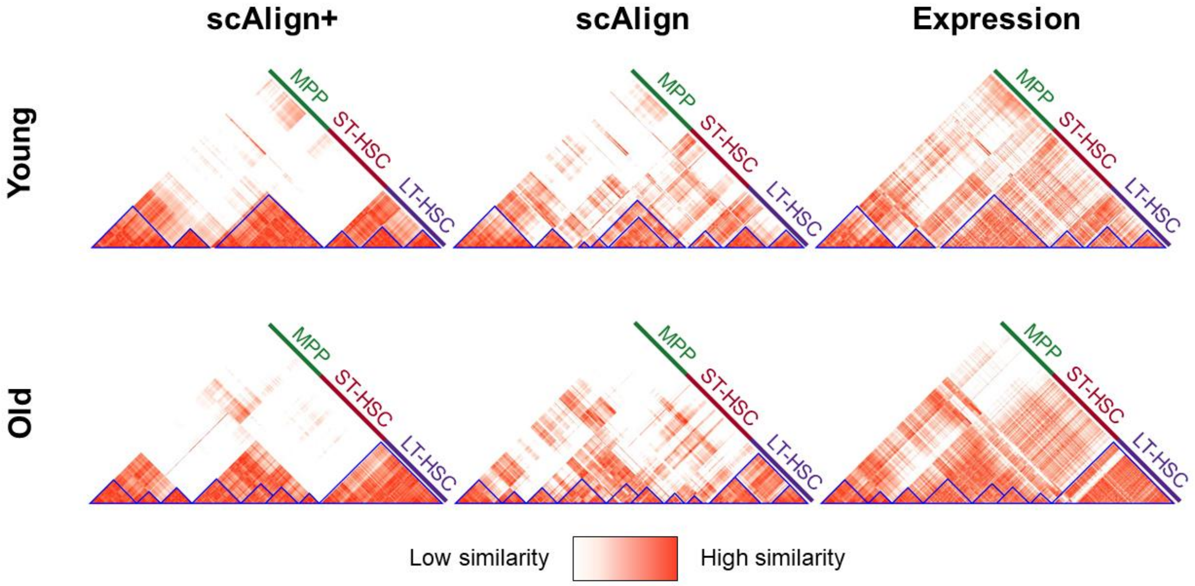

**Supplementary Figure 9: Cell-cell similarity matrix after supervised or unsupervised training of scAlign compared to the gene expression-based similarity matrix.** Heatmaps of cell-cell similarity illustrate the agreement between training scAlign+ with all cell type labels (supervised) or scAlign without any cell type labels (unsupervised) for both the young and old cells in Kowalczyk et al., or when directly measuring similarity in gene expression space. Clusters of cells are highlighted within and across each heatmap by blue triangles.

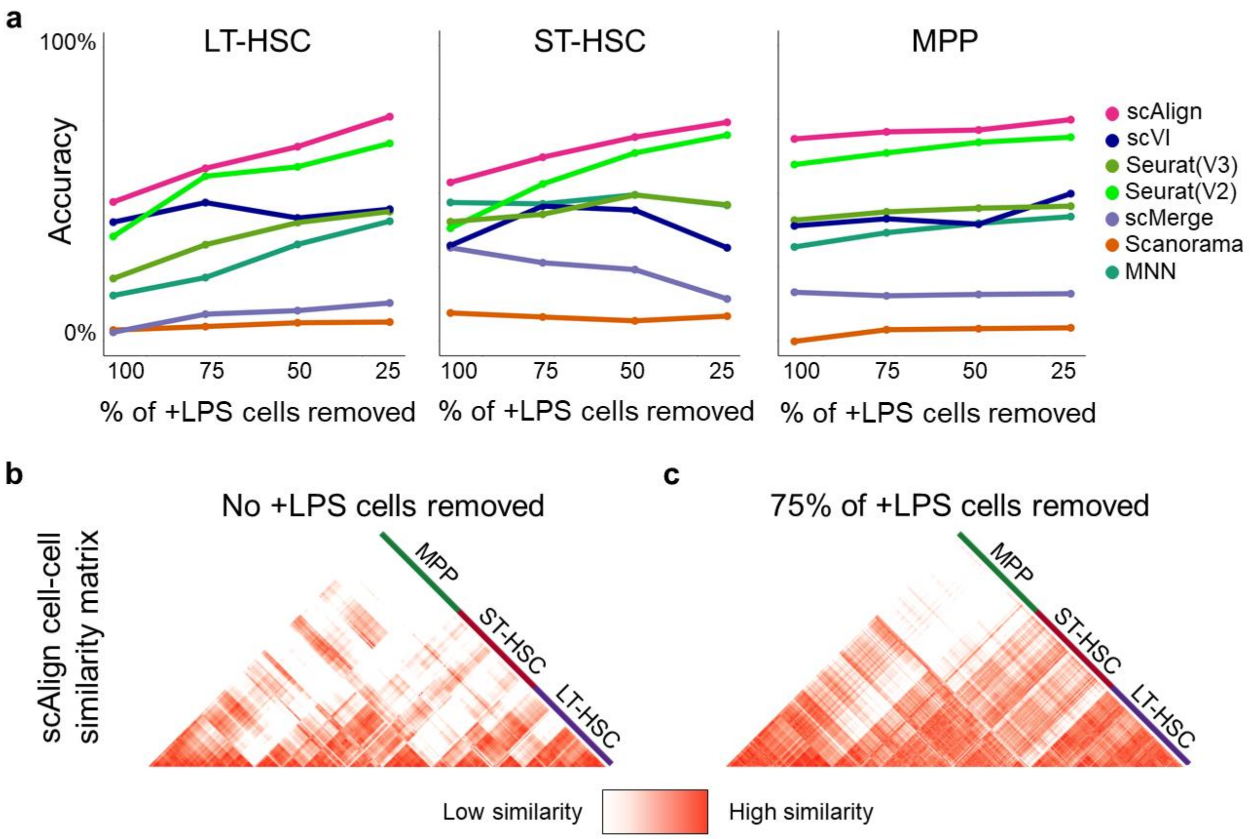

**Supplementary Figure 10: Performance of alignment methods on the Mann et al. benchmark after removing cells**. (**a**) Accuracy of classifiers on the Mann et al. benchmark, when removing a percentage of either LT-HSC, ST-HSC or MPP cells from the old condition. scAlign outperforms all other methods robustly on the full dataset, and most methods perform worse as more cells are removed from the old condition. (**b**) Heatmap showing the pairwise similarity matrix for the stimulated (+LPS) cells from Mann et al. when no cells have been removed. (**c**) Heatmap showing the pairwise similarity matrix for the stimulated (+LPS) cells from Mann et al. after keeping only 75% of the target cells from all cell types.

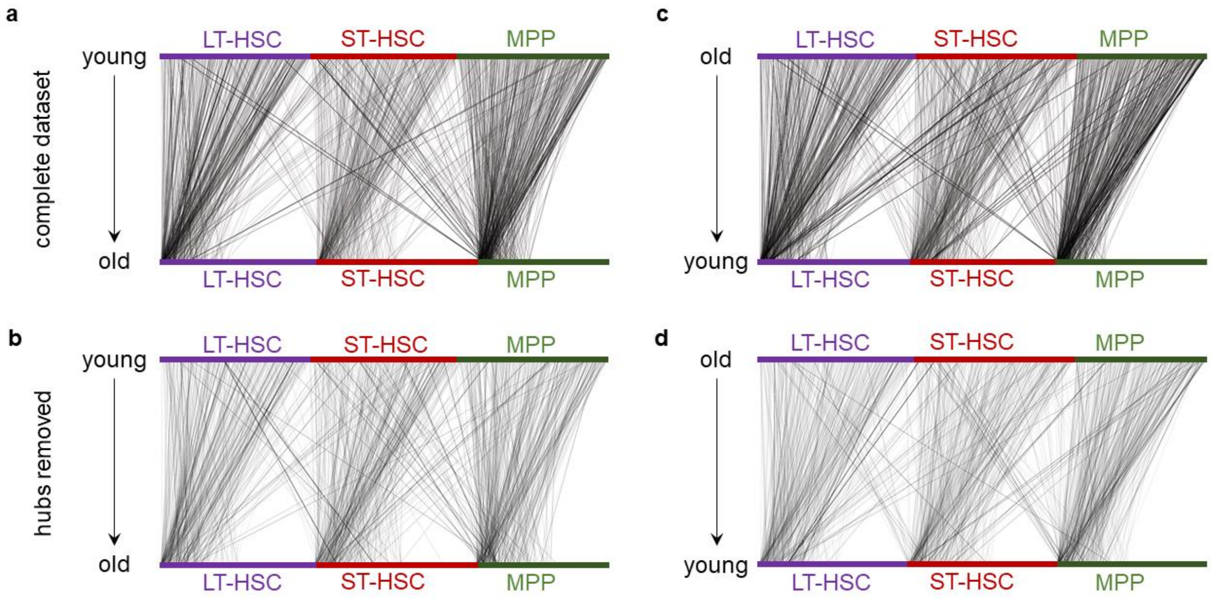

**Supplementary Figure 11: Random walk probabilities measured before and after removing cells frequently visited on random walks during scAlign training on Kowalczyk et al.** (**a**) Top layer of nodes represent cells in the young condition, while the bottom layer of nodes represents cells in the old condition. Nodes are sorted by degree such that hubs (cells visited frequently on the random walks) are grouped on the left of each cell type in the old condition. An edge represents a high probability walk from a young cell to an old cell, where thicker edges indicate more frequent walks. (**b**) Same as (a), but after removing the hubs identified by selecting nodes with degree above the 90$^{th}$ quantile. (**c**) Same as (a), but the top layer now represents old cells, while the bottom layer represents young cells, and edges represent random walks from old to young cells. (**d**) Same as (c), but after removing the hubs.

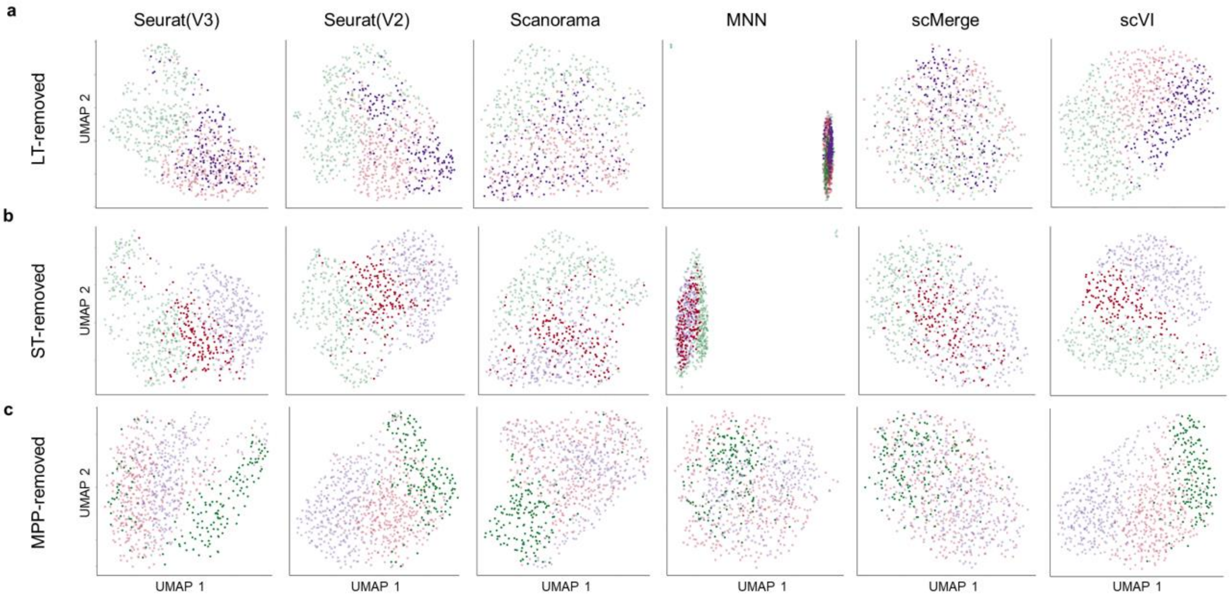

**Supplementary Figure 12: Comparison of alignment methods when cell types are removed from the Kowalczyk et al. benchmark. (a)** t-SNE visualization of alignments after removing LT-HSCs from the stimulated cells. The control LT-HSCs are highlighted to show the amount of incorrect overlap with either the ST-HSC or MPP cells. **(b-c)** Similar to (a), but with ST-HSCs or MPPs removed, respectively.

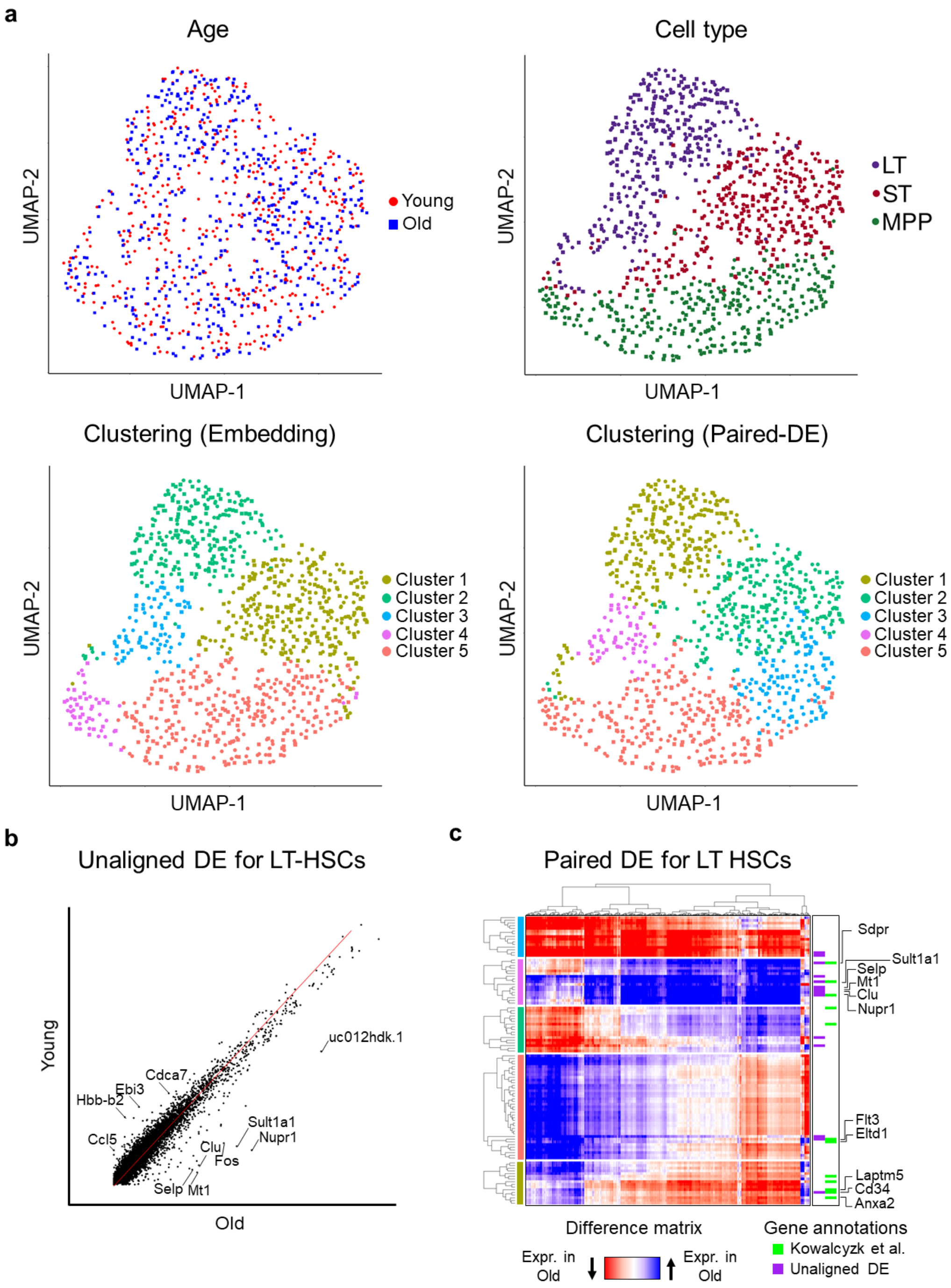

**Supplementary Figure 13: Alignment of Kowalczyk et al. identifies subpopulations of LT-HSC(s) with unique response to age.** (**a**) UMAP visualization of young and old mouse cells after alignment by scAlign. Each cell is colored by age, cell type, clustering in alignment space or clustering on scAlign's state variance map. (**b**) Scatterplot of significantly differentially expressed genes identified by comparing cluster averages. (**c**) Heatmap of the state variance map (paired difference for each cell interpolated into the young and old condition). Each gene is annotated based on the differential expression results from Kowalczyk et al. and comparison of cluster averages.

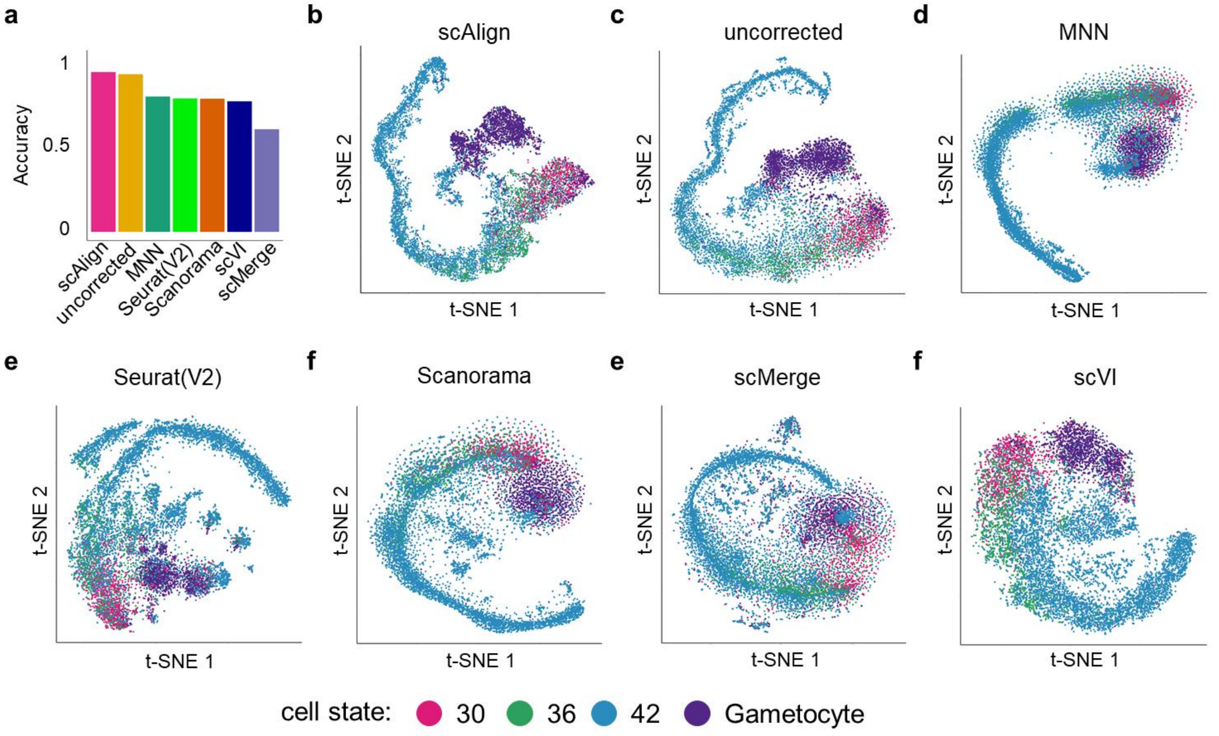

**Supplementary Figure 14: Alignment of +/-Shld parasites**. (**a**) Accuracy of different alignment methods after aligning +/-Shld parasites, where accuracy is based on the notion that gametocyte cells from the +Shld condition should not be aligned to any parasites in the -Shld condition. (**b-f**) tSNE visualizations of +/-Shld parasites aligned together, using different alignment methods.

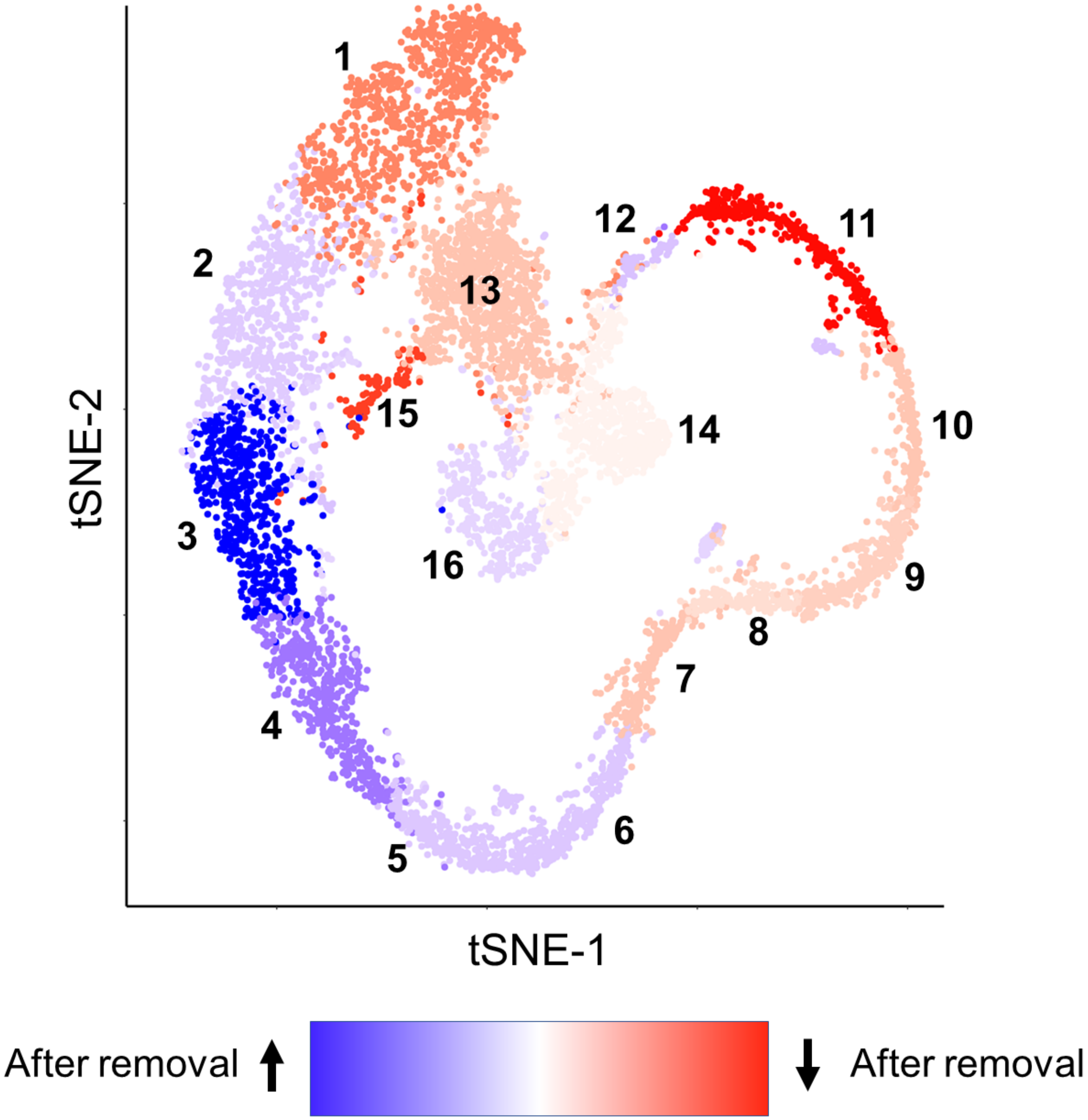

**Supplementary Figure 15: Removal of -Shld parasites from gametocyte clusters leads to more diffuse round trip probabilities from cluster 13.** The difference heatmap for round trip probabilities for parasites in cluster 13 (rows) in the presence of cells from -Shld grouping with +Shld gametocytes and after removal of the -Shld cells grouping with +Shld gametocytes. Dark blue indicates an increase in round trip probability after removal and red indicates a decrease in round trip probability after removal.

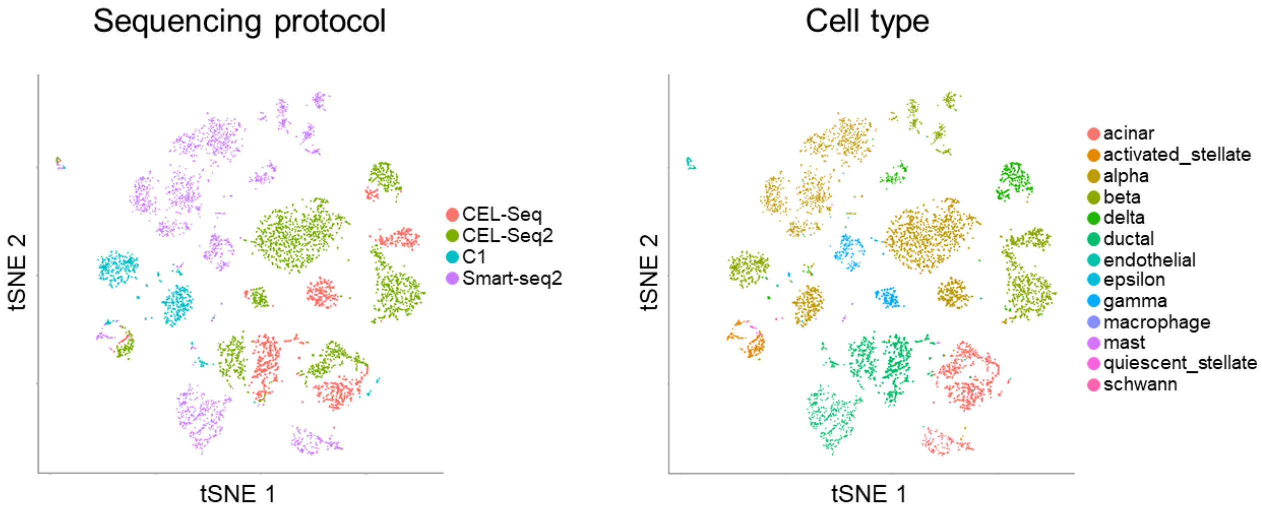

**Supplementary Figure 16: Unaligned pancreatic datasets exhibit protocol specific effect**. tSNE visualization of the unaligned highly variable genes, cells are colored by sequencing protocol and cell type.

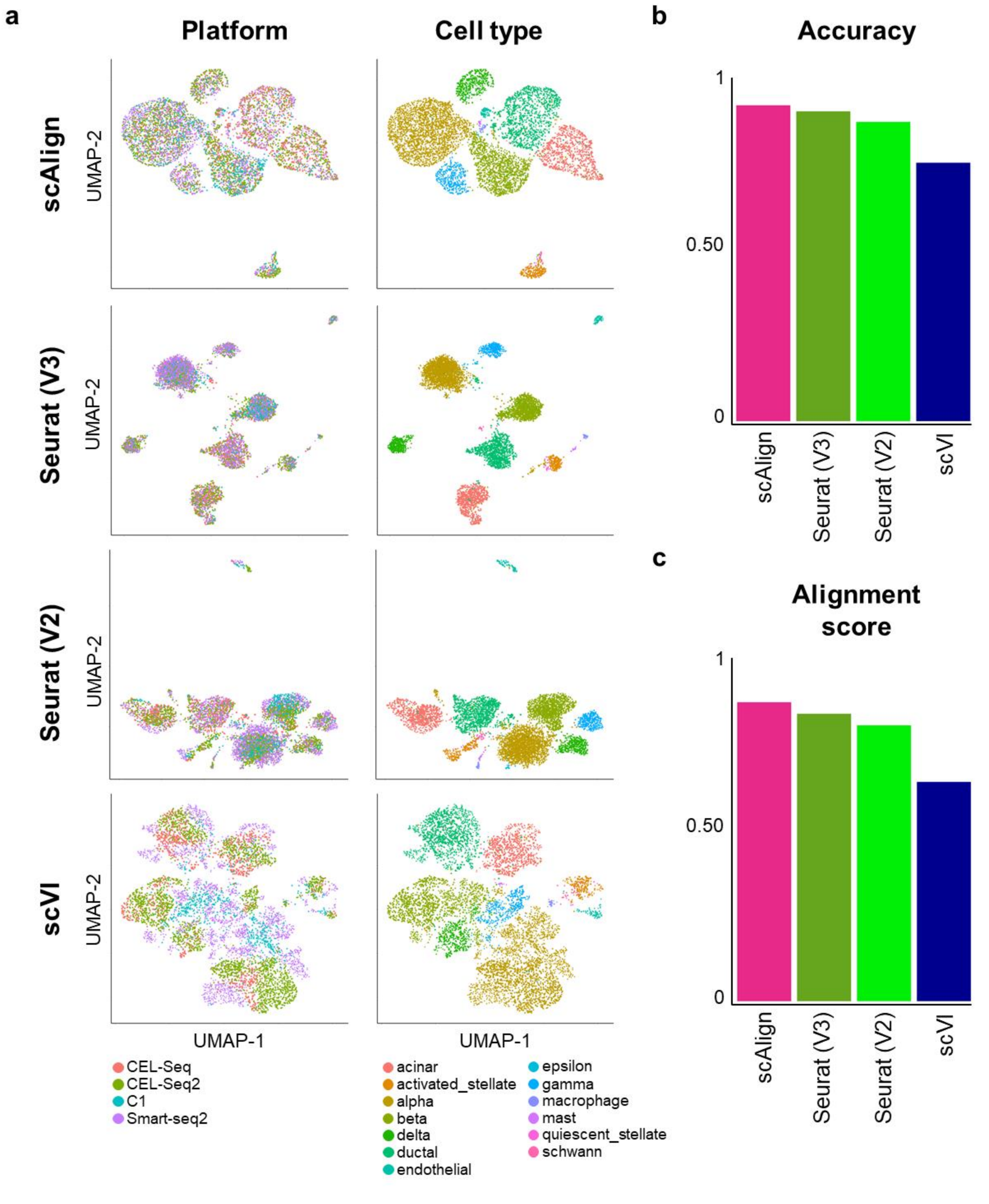

**Supplementary Figure 17: Alignment of all four pancreatic islet datasets**. (**a**) UMAP visualization of the aligned embeddings produced by scAlign colored by platform or cell type. (**b**) Barplot visualizing the composite accuracy for each method. (**c**) Barplot visualizing the alignment scores for each method.

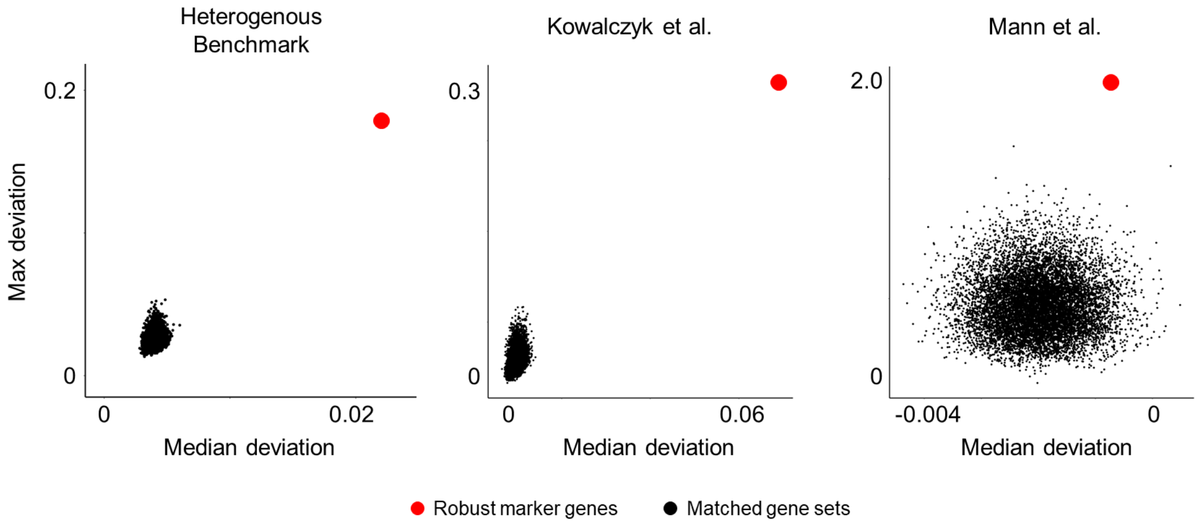

**Supplementary Figure 18: Robust cell type marker genes drive alignment**. For the set of cell type marker genes robustly identified across conditions, we perturbed their expression in cells and measured the corresponding deviation in cell state space embeddings. We repeated this experiment for gene sets matched for size and relative expression. Perturbation of the robust cell type marker genes led to systematically larger deviations in cell state embeddings compared to control gene sets, indicating that robust cell type marker genes contribute more to alignment than expected by chance.

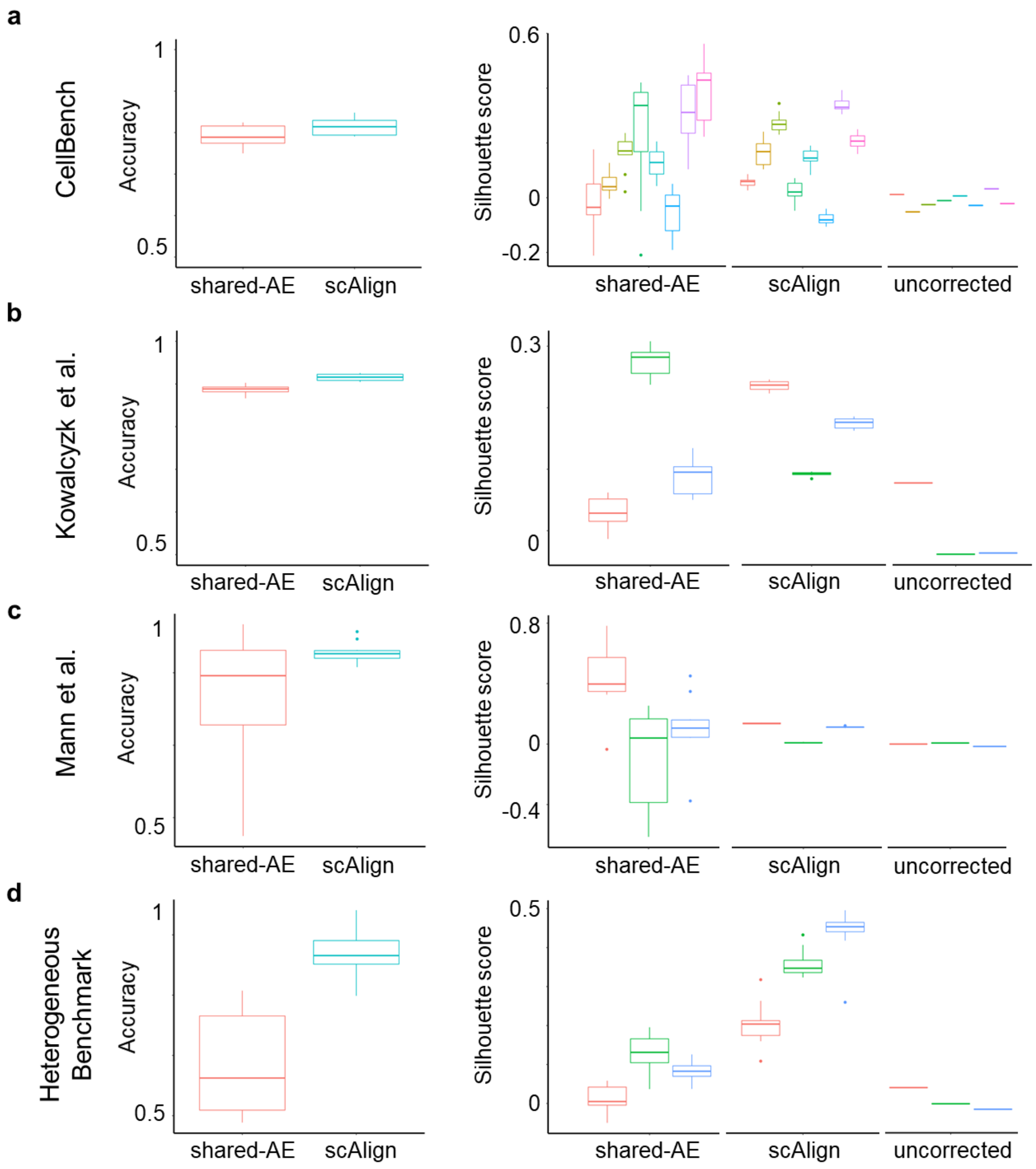

**Supplementary Figure 19: scAlign outperforms a shared autoencoder (shared-AE) with similar network architecture, in terms of alignment accuracy and maintaining fidelity of the cell-cell similarity matrix**. (**a-d**) Box and whisker plots of alignment quality metric after 10-fold CV for both the shared autoencoder and scAlign. Silhouette score is illustrated for unaligned data, cell embeddings for scAlign and for the shared autoencoder.

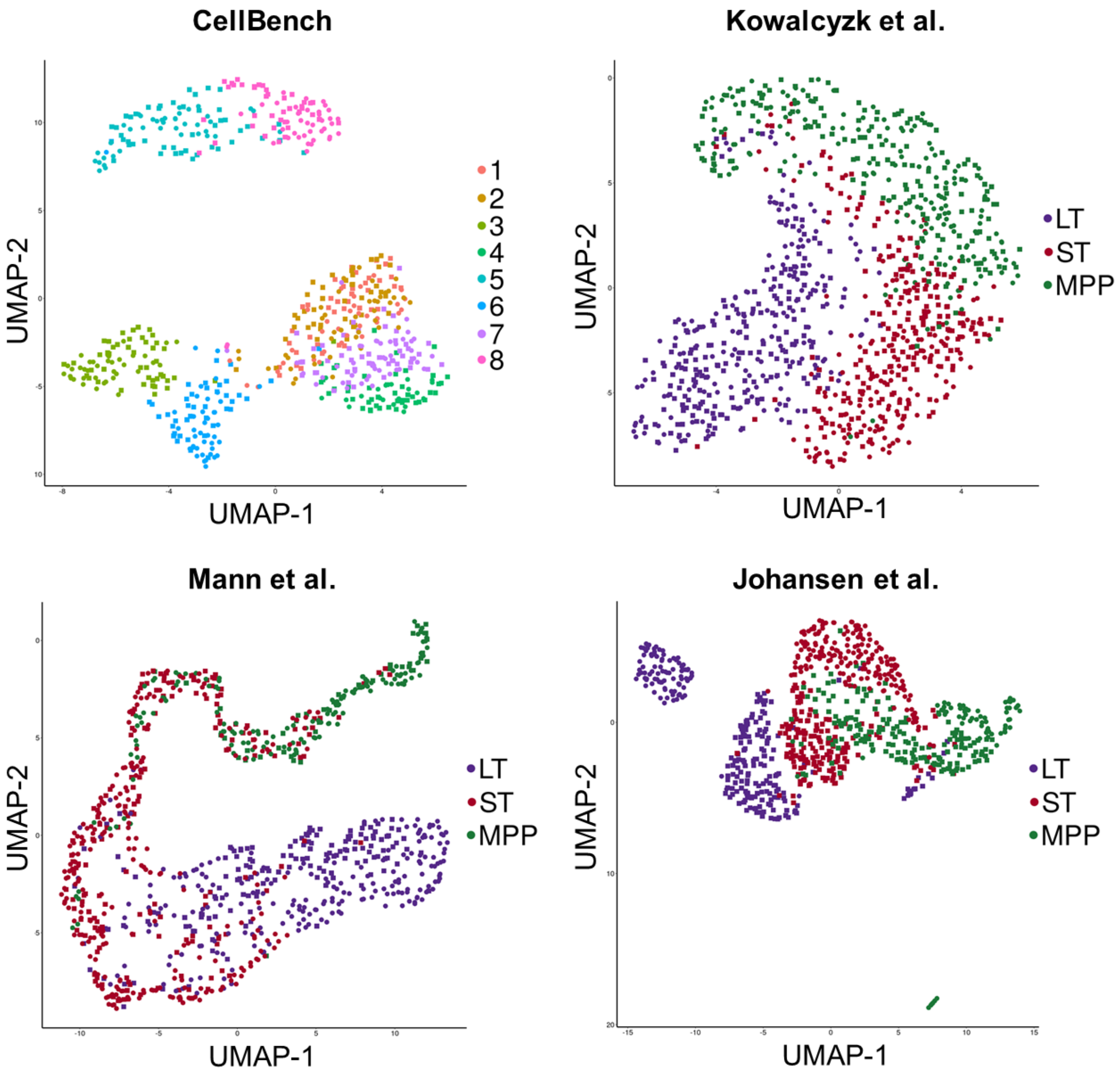

**Supplementary Figure 20: Comparison of shared autoencoder cell embeddings after alignment of the four benchmark datasets.** UMAP visualization shows the cell embeddings colored by cell type for each of the four benchmark datasets.

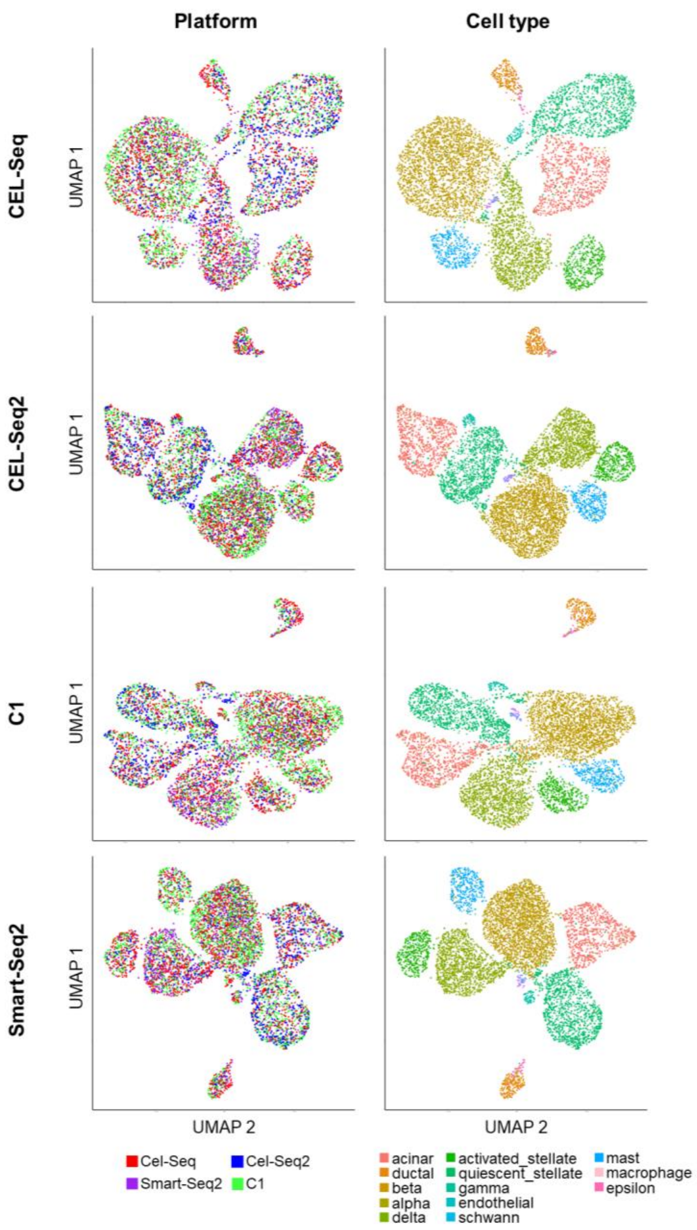

**Supplementary Figure 21: Comparison of scAlign alignment of pancreatic islet cells using each protocol as a reference.** UMAP visualizations after alignment where a single protocol is used as a reference (y-axis) and colored by both platform and cell type (x-axis) annotations.

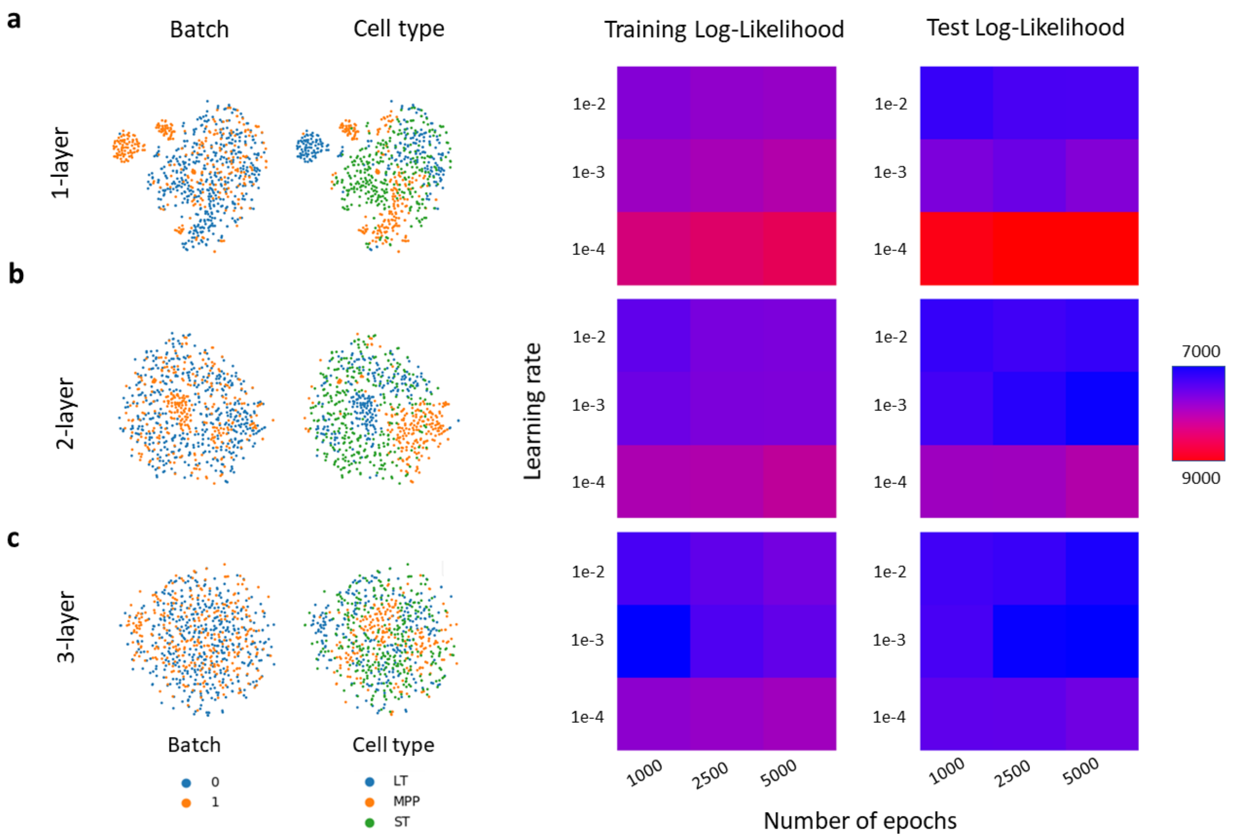

**Supplementary Figure 22: scVI grid parameter search procedure identifies optimal parameterization.** (**a**) (left) tSNE visualization shows the latent dimensions inferred by scVI, colored by batch (condition) and cell type, after alignment using the optimal parameters identified by grid search. (right) Parameter search results for both the training and test set with respect to log likelihood, where the x-axis is the learning rate and y-axis is the number of epochs for the respective number of network layers. (**b-c**) Same as (a), but for different numbers of hidden layers in the network.
